# DMcloud: Macromolecular Structure Modeling Using Local Structure Fitting for Medium to Low Resolution cryo-EM maps

**DOI:** 10.64898/2026.06.12.731990

**Authors:** Genki Terashi, Xiao Wang, Yuanyuan Zhang, Han Zhu, Joon Hong Park, Daisuke Kihara

## Abstract

Cryogenic electron microscopy (cryo-EM) has become an essential experimental approach in structural biology for determining macromolecular structures. When the resolution of a cryo-EM map is worse than approximately 5 Å, fitting known or predicted molecular models into the map becomes a common strategy for interpretation. However, accurately fitting biomolecular models into cryo-EM maps, particularly for large macromolecular complexes, remains challenging when the input structure models contain errors or are in a conformation different from that represented in the map. Here, we present DMcloud, a method for local structure fitting of proteins and nucleic acids in cryo-EM maps. Instead of forcing an entire input model into the map, DMcloud divides input structures into local regions, identifies regions that are supported by the density, removes unsupported regions, and assembles the retained regions into a final model. We benchmarked DMcloud on 176 cryo-EM maps, including intermediate and high-resolution maps that include proteins, DNAs, or RNAs. For EM maps in the 5.0-10.0 Å and 2.5-5.0 Å resolution ranges, DMcloud achieved average sequence modeling coverage of 0.49 and 0.70, respectively. For DNA/RNA maps, DMcloud achieved an average sequence coverage of 0.75. Across all datasets, DMcloud consistently outperformed existing methods in model accuracy, map-model correlation, and modeling coverage.

## Introduction

Cryo-electron microscopy (cryo-EM) has revolutionized structural biology by enabling the determination of protein and nucleic acid macromolecular structures that were previously difficult or impossible to resolve using conventional techniques such as X-ray crystallography^1,2^. However, interpreting and modeling biomolecular structures from cryo-EM maps, particularly those with resolutions worse than approximately 5 Å, remains a significant challenge^3^. Existing modeling methods often struggle to accurately trace protein and DNA/RNA backbone structures in such lower-resolution maps. Even in maps resolved at around 2.5 to 4.0 Å, modeling errors have been reported, including incorrect assignments of amino acid types^4,5^. When the resolution of a cryo-EM map is worse than 5 Å, employing a model fitting approach, rather than a de novo main-chain tracing method^6^, is often more practical choice^7^. Structures to be fit into the map are typically obtained from the Protein Data Bank (PDB)^8^ or predicted using structure prediction methods such as AlphaFold2 (AF2)^9^. These models are then fit to the map using rigid-body fitting methods^10–13^.

Several model-map fitting methods have been proposed. DEMO-EM2^14^, for example, employs an iterative assembly process that integrates both chain-level and domain-level alignment of AF2-predicted models with Fast Fourier Transform (FFT)-based fitting^15^. EMBuild^16^ fits AF2 models using a combination of rigid-body global fitting, domain-based semi-flexible refinement, and graph-based iterative assembly of chains. EMBuild operates on predicted residue positions derived from the map rather than the raw density map itself. Our group developed DiffModeler^17^, which uses a diffusion model^18^, a type of generative deep learning model, to enhance main-chain positions in cryo-EM maps, thereby facilitating more accurate fitting of AF2-predicted structures.

Although conceptually straightforward, the fitting approach faces three major challenges that can reduce modeling accuracy. The first challenge concerns the accuracy of the input structures. Even experimentally determined structures of individual proteins may adopt conformations that differ from those present in the cryo-EM map, leading to potential misfitting. When a predicted structure by AF2 is used, it is often the case that domain of a protein is incorrect, despite that individual domains or parts in a structure are correct. When a structure predicted by AF2 is used, it is often the case that the relative domain orientation of the protein is incorrect, even though the individual domains or local regions are accurately modeled^19^. The second challenge lies in the low-resolution nature of the map. As resolution decreases, salient structural features become less distinct, making it more difficult to fit a structure in the correct orientation. Additionally, a third challenge is that current methods often require explicit chain information, such as the number of chains for each component, which may not be available prior to full structural determination by cryo-EM.

To address these challenges, here we developed DMcloud, a biomolecular local structure fitting method that combines point cloud matching techniques for local structure fitting and a structure-enhanced EM map generated using a diffusion model. DMcloud performs local structure fitting using two key computational techniques: RANSAC^20^ (Random Sample Consensus) for robust initial alignment and DBSCAN^21^ (Density-Based Spatial Clustering of Applications with Noise) for identifying matching local structure regions and refining the alignment. This approach enables more accurate assembly of AF2 models with cryo-EM maps by mitigating inaccuracies in domain orientations and other modeling errors present in AF2 models. Notably, DMcloud selectively fits only the local regions of AF2 models that are compatible with the observed map density.

We benchmarked DMcloud on three datasets: 71 intermediate resolution maps (5 - 10 Å), 50 high resolution maps (2.5 - 5.0 Å), and 55 maps containing both DNA/RNA and proteins. Across all datasets, DMcloud consistently outperformed four existing methods^14,16,17,22^ in terms of model accuracy, map-model correlation, and modeling coverage. DMcloud is available as open-source software and through a user-friendly web server, facilitating broad accessibility for structural biologists.

## Results

### Overview of the DMcloud protocol

DMcloud performs local structure fitting of models into a cryo-EM map in four main steps (Fig. 1). The required inputs are a cryo-EM map and a set of unique protein structure models. If a protein complex contains multiple copies of the same chain, only a single instance of that structure is needed; users are not required to specify the number of copies. DMcloud automatically identifies all fitting locations of each chain structure within the map. This is a notable advantage over existing methods, which typically require users to explicitly provide the number of copies of each chain in the complex.

**Figure 1.**
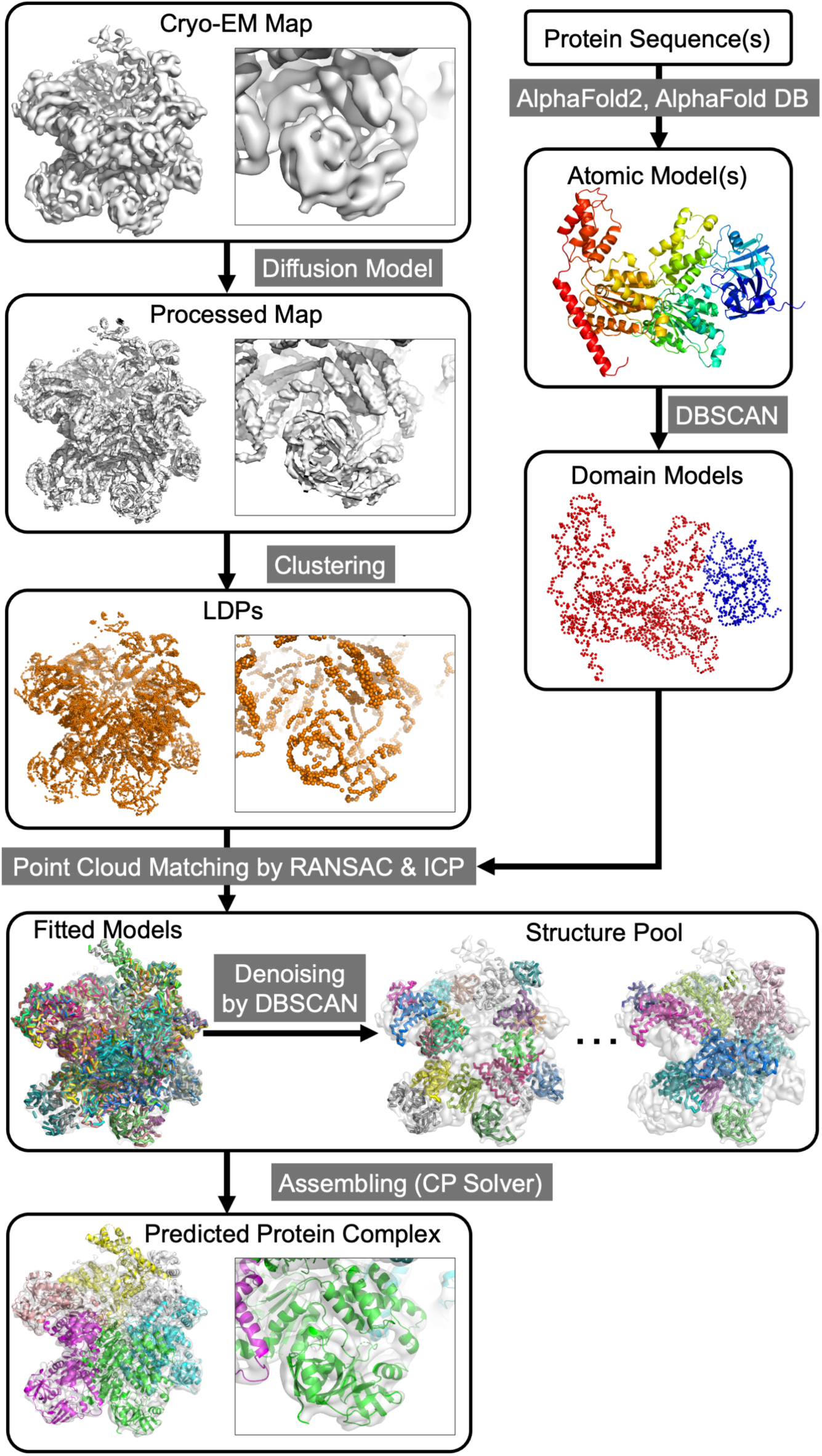
Overview of DMcloud. Main steps of structure fitting procedure are illustrated. A cryo-EM map of ADP-bound VAT complex (PDB ID 5G4F^26^, EMD-3436; resolution 7.0 Å) and an AF2 model are used as an example. The inset shows the N-terminal domain, residues 1-176. In the left branch, the input cryo-EM map is processed using a diffusion model to enhance backbone visibility and converted into point cloud data with local dense points (LDPs). In the right branch, the AF2 model is represented as a point cloud data with Cα positions and split into domains using DBSCAN. The map and each domain are aligned and refined. This step produces a pool of fitted local structures in the map. Finally, fitted domain structures are assembled into the final protein complex.

The protocol begins by processing the input cryo-EM map using a diffusion model to enhance backbone visibility. Next, representative high-density points in the map, referred to as local dense points^23^ (LDPs) are identified. These LDPs serve to convert the map into a point cloud representation. Meanwhile, the input protein structure models, which can be those by AF2, are also represented by point clouds with Cα positions, and split into domains using a DBSCAN. Thus, both map and structure (domain) models are converted into point clouds. Subsequently, point clouds of the domain models are aligned to the LDPs of the map using RANSAC, followed by refinement of alignment through Iterative Closest Point^24^ (ICP) and denoising by DBSCAN. The fitted structures are then pooled and assembled using a constraint programming (CP) solver^25^ to model the final protein complex. In the following, we explain each step. Further information can be found in the Methods section.

### Enhancement of backbone positions in the cryo-EM map using a diffusion model

Accurate fitting of structure models is challenging in intermediate-resolution cryo-EM maps. To improve fitting accuracy, DMcloud enhances the input map using a diffusion model^27^ developed for DiffModeler^17^. We did not newly train the diffusion model in this work. The diffusion model emphasizes regions corresponding to the protein backbone. Specifically, the cryo-EM map is scanned using 64³ Å³ boxes at 1 Å intervals. Within each box, the predicted backbone positions are enhanced by increasing the local density, while non-backbone regions are attenuated. This preprocessing step facilitates more accurate fitting of structure models into the map.

### Representing the modified map as a point cloud

Subsequently, the modified map by the diffusion model is represented by point cloud of LDPs. LDPs are representative points of high-density regions, which are likely to correspond to atoms in proteins. Identification of LDPs is performed with the mean-shift algorithm^23^, which is a local classification method. Typically, LDPs are placed along the backbone of proteins with additional LDPs at side-chain densities. On average there are 3 to 4 LDPs for a residue in a protein.

### Protein domain identification and point cloud representation

The structural model of protein chains to fit in the map are obtained from the AlphaFold Database^28^. For homo-oligomeric proteins or proteins containing homo-oligomeric components, DMcloud requires only unique single-chain models. A key feature of DMcloud is its ability to automatically identify the number and spatial locations of these single-chain models present within the input cryo-EM map.

Protein models are first segmented into domains by clustering main-chain atoms (Cα, N, and C) using DBSCAN. The clustering takes into account both the interatomic distances and the pLDDT confidence scores from the AF2 models, which are used as weights. As a result, each domain is represented as a point cloud of main-chain atoms.

### Matching protein domains with the map

Next, protein model domains are matched and aligned to the map by a point cloud matching procedure. Similar to the domain segmentation applied to the protein models, the map is also segmented into local regions, each consisting of LDPs within a 15 Å radius. Each point in the point clouds of both the map and the protein domains is characterized using a Fast Point Feature Histograms (FPFH)^29^, which capture local geometric features of the surrounding region. The alignment is computed exhaustively between every pair of the protein domains and the segmented map regions and ones that meet a predefined similarity threshold are retained.

Alignment of a protein domain and a local map region is initially performed using RANSAC followed by refinement with ICP. RANSAC begins by aligning three points that have similar FPFH descriptors and includes additional points that fall within 3 Å of their counterparts. This initial alignment is then refined by ICP, which iteratively improves the fit by incorporating more corresponding points. Finally, DBSCAN is applied to the refined alignments to identify consistent clusters and remove outliers. For each cluster, the resulting alignment is recorded as a fitted model in the structure pool. Since RANSAC performs alignments from various seed points, multiple distinct alignments may be generated and retained as candidate fits. Typically, for a single protein domain, approximately 140 alignments to the map are stored. This is especially relevant for homomultimeric complexes, where a single domain structure can correctly fit into multiple equivalent positions within the map.

### Assembling fitted structure domains

The previous matching step will produce about 100-1000 fitted structure models. These fitted models are finally combined and assembled by combinatorial optimization using a constraint programming (CP) solver^30^. The combination optimizes a score that consists of a score that considers the fitness of the protein domain structure to LDPs in the map using the local distance difference test (LDDT) score^31^ and a penalty term that considers clashes between overlapping domains.

### Fitting nucleic acid models

DMcloud extended its predecessor, DiffModeler, by enabling structure fitting of DNA/RNA structure models. There are several key differences between nucleotide fitting and protein fitting. When processing cryo-EM maps containing DNA or RNA, DMcloud employs a deep learning model with a U-Net++ architecture, originally developed for CryoREAD^32^ to detect the locations of phosphate atoms and sugar groups of nucleic acids, instead of using a diffusion model of cryo-EM maps. LDPs are then calculated based on these detected sites, with classifications according to their atom types. DMcloud performs two-color point cloud matching, distinguishing between phosphate and sugar points to more robustly fit nucleotide structures. DMcloud uses DNA/RNA structure models deposited in the PDB, which are identified through sequence searches for the cases presented in this study. For targets that contain both nucleic acids and proteins, we first constructed a DNA/RNA model and then performed the DMcloud protocol to fit proteins to the map.

### Structure modeling results of DMcloud

We tested DMcloud on four different datasets of cryo-EM maps. The first dataset is 71 maps determined at an intermediate resolution (5.0 to 10.0 Å). Next, we also tested the method on 50 higher resolution maps of 2.0 to 5.0 Å. Additionally, we tested on 55 maps that contain DNA/RNA and proteins. Proteins in these datasets are nonredundant to the training and validation datasets used for developing the diffusion model in the DiffModeler, which we also used in DMcloud.

### Modeling performance on intermediate resolution cryo-EM maps

First, we benchmarked DMcloud on the same dataset as used in the DiffModeler paper^17^ so that we can compare the performance between them. It consists of 71 maps at resolutions between 5.0 to 10.0 Å, which is the primary target resolution range for this structure fitting method (Supplementary Table S1). The corresponding PDB structures to the maps contain 1 to 62 protein chains, which have 814 to 18,080 residues in total. The same modified maps using the diffusion model and the same AlphaFold2 models (Supplementary Table 2). We compared the performance of DMcloud against four existing methods: DiffModeler, EMBuild, and DEMO-EM2, which are structure-fitting approaches, and ModelAngelo^22^, a de novo structure-building method. For the three fitting-based methods, we used the same AF2 models as those employed by DMcloud to ensure a fair comparison.

In Fig. 2a, the methods were evaluated based on sequence coverage, defined as the fraction of residues in the model placed within 3 Å of the reference structure. DMcloud achieved higher sequence coverage than all other methods for nearly all targets, except for a few targets where EMBuild or DiffModeler had a higher value. The average sequence coverage of DMcloud was 0.491, followed by DiffModeler (0.419) and EMBuild (0.412). DEMO-EM2 showed a lower average value of 0.201. This is not the resolution range where the de novo modeling method, ModelAngelo is designed for (0.004). In Fig. 2b, we further examined the models in terms of cross-correlation, which quantifies the overall global fit of the structure model to the map (CC_box with *phenix.map_model_cc* in Phenix^33^). As shown in the plot, DMcloud showed a higher cross-correlation for almost all the maps than the other methods. The average CC_box for DMcloud was 0.669, outperforming DiffModeler (0.587), EMBuild (0.587), DEMO-EM2 (0.561), and ModelAngelo (0.163). Backbone F1 score shown in Fig. 2c evaluates both the residue (or nucleotide for DNA/RNA) position accuracy and the completeness of the predicted model relative to the reference PDB structure. Consistent with the previous two plots (Fig. 2a, 2b), DMcloud overall clearly shows higher values for almost all the cases compared to the other methods. It is notable that DMcloud achieved consistently higher evaluation scores than DiffModeler. Since the same diffused maps and AF2 models were used for both, the performance difference arises solely from the newly developed local fitting procedure in DMcloud. In Extended Data Figure 1, we additionally analyzed the modeling performance using other similar evaluation metrics. Individual data is provided in Supplementary Table S3.

**Figure 2.**
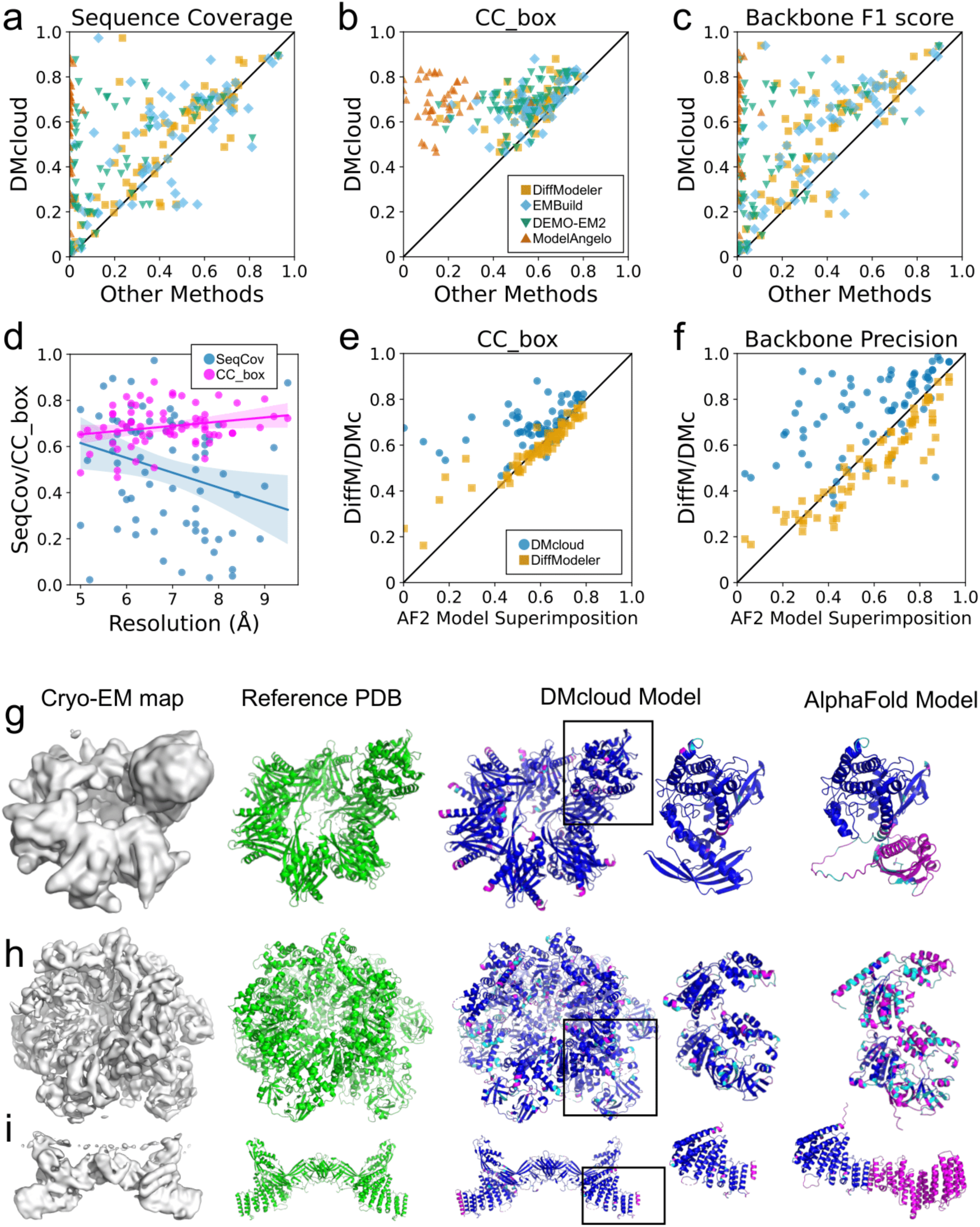
Modeling results on the intermediate-resolution dataset. In panel **a, b, c**, the performance of DMcloud (y-axis) was compared with four other methods (x-axis), DiffModeler (squares in yellow), EMBuild (diamonds in cyan), DEMO-EM2 (downward triangles in blue), and ModelAngelo (upward triangles in brown). **a.** Sequence Coverage, the fraction of correctly placed residues; **b.** CC_box, cross-correlation of the model and the cryo-EM map; **c.** Backbone F1 score, balanced precision and recall in model accuracy. Source data is provided in Supplementary Table S3. **d.** Sequence Coverage (Blue) and CC_box (magenta) of models by DMcloud relative to the map resolution. The lines represent the linear regression line. Sequence Coverage shows a negative correlation with resolution (R = 0.276). CC_box shows a positive correlation (R = 0.224). The shaded regions indicate the 95% confidence intervals of the regression lines. **e.** CC_box of models generated by DMcloud and DiffModeler relative to CC_box of AF2 model superimposition. AF2 models were superimposed onto the reference PDB structures using structural alignment with PyMol’s *align* command. **f.** Backbone precision of models generated by DMcloud and DiffModeler relative to the backbone precision of AF2 model superimposition. Panel **g-j**. Rows from left to right are: (i) the surface representation of the EM density map at the author-recommended contour level; (ii) the reference PDB structure; (iii) comparison between the DMcloud model and the reference PDB structure, both the entire complex structure and the individual chain; (iv) and a comparison between the AlphaFold2 model and the reference PDB structure. In the fourth and fifth columns, residues in the structure models are color-coded: blue indicates residues correctly placed within 3 Å of their corresponding residues in the PDB structure; cyan marks residues positioned within 3 Å of some residue in the PDB structure but not their corresponding amino acid; and magenta highlights residues placed more than 3 Å away from any residue in the reference PDB structure. **g.** Calcium/calmodulin-dependent protein kinase II subunit alpha (EMD-21536; resolution: 6.6 Å; PDB ID: 6W4P). Sequence Coverage 0.973, CC_box 0.880; Backbone F1 score 0.939. In the DMcloud model panel, a hub protein (chain B) and a kinase domain (chain H on the top of the complex) are marked with a box and magnified. On the right panel, the AF2 model of the single-chain, which consists of a hub and a kinase domain, is shown. **h.** ADP-bound Valosin-containing protein-like ATPase (EMD-3436; res. 7.0 Å; PDB ID: 5G4F). Sequence Coverage 0.692; CC_box 0.832; Backbone F1 score 0.728. In the DMcloud model panel, chain A is marked with a box and magnified. In the AF2 model panel, chain A is shown. **i.** Metazoan retromer and the sorting nexins3 complex (EMD-12221; res. 9.5 Å; PDB ID: 6VAC). Sequence Coverage 0.876; CC_box 0.722; and Backbone F1 score 0.878. In the DMcloud model panel, chain A is marked with a box and magnified. In the AF2 model panel, chain A is shown.

In the next three panels, we analyzed how DMcloud’s modeling quality is influenced by two key factors: map resolution and AF2 model accuracy. Figure 2d shows model accuracy, measured by sequence coverage (blue) and CC_box values (magenta), as a function of map resolution. Sequence coverage exhibits a slight decrease as resolution worsens. At map resolutions of 5–6 Å, the average sequence coverage was 0.56, which decreased to 0.38 at resolutions of 8 Å or worse. In contrast, CC_box remains stable across resolutions, suggesting that while the overall structures are still captured at low resolutions, the accuracy of detailed residue positions is affected. In Figure 2e and 2f, modeling performance of DMcloud and DiffModeler were compared relative to AF2 model quality. To quantify AF2 model accuracy, we superimposed AF2 models to the PDB entry and computed the fraction of model residues that are within 3 Å to the corresponding residues in PDB using phnix.chain. For both CC_box (Fig. 2e) and backbone precision (Fig. 2f), DiffModeler shows a strong dependence on AF2 model quality, as it relies on global superimposition of AF2 models. In contrast, DMcloud performs local structure fitting by identifying compatible regions in AF2 models with the map, thereby reducing false positive placements and producing models that fit the input cryo-EM maps far more accurately.

### Examples of structure modeling for intermediate-resolution maps

Figures 2g–2i present three examples of structure models built by DMcloud. Extended Data Figure 2 shows corresponding structure models by the other three methods. In the right panels, residues in the predicted models are colored blue, cyan, or magenta to indicate positional accuracy: blue for residues correctly placed within 3 Å of the corresponding position; cyan for residues located in the correct protein region but not within 3 Å of the correct position; and magenta for residues more than 3 Å from any residue in the reference PDB structure. The rightmost panel shows the individual AF2 model fitted to the map using the same color scheme.

The first example (Figure 2g) is calcium/calmodulin-dependent protein kinase II subunit alpha (CaMKII-alpha), modeled from a 6.6 Å EM map (EMD-21536). This protein complex is a 13-mer consisting of 12 hub domains and a kinase domain, totaling 1,789 residues (PDB ID: 6W4P^34^). The chain in the PDB entry, which includes the ring-shaped 12 hubs and the kinase domain at the top of the ring, is nearly identical to the AF2 prediction (99.6% identity, differing by only one residue); therefore, only one AF2 conformation was considered (shown in the rightmost panel). A major modeling challenge was that either the hub domain (Asp340–His478) or the kinase domain (Met1–Arg311) must be correctly extracted from the AF2 model and locally fitted to the map, as the domain orientations in the AF2 prediction do not match the conformation observed in the EM density. When aligning the AF2 model to the hub domain in the reference PDB structure, only 285 out of 478 residues in the AF2 model can be correctly placed (the right panel). Due to this, DiffModeler, which performs global fitting, had a limited performance with sequence coverage of 0.235, CC_box of 0.677, backbone F1 score of 0.105. In contrast, DMcloud accurately fit 12 hub domains and a single kinase domain into the map, achieving the highest sequence coverage (0.973), CC_box (0.880), and Backbone F1 score (0.939) compared to other methods (Extended Data Figure 2a).

The second example (Figure 2h) is the ADP-bound Valosin-containing protein-like ATPase of *Thermoplasma acidophilum* (VAT) complex, modeled from a 7.0 Å map (EMD-3436; PDB ID: 5G4F^26^). It is a homo hexamer with 4,356 residues. The VAT has different conformational states, and the AF2 predicted an open conformation that differed from the reference PDB structure. When directly aligning the AF2 model to the reference PDB structure, the AF2 model could only place 170 out of 745 residues of a subunit correctly (Figure 2h, right panel). Due to the different conformation of the AF2 model, the global fitting by DiffModeler only achieved sequence coverage of 0.355, CC_box of 0.652, and backbone F1 score of 0.350 (Extended Data Figure 2b). In contrast, DMcloud adjusted the orientation of the domains of the AF2 model to form the closed conformation when it was fitted to the map, achieving overall correct model with a sequence coverage of 0.692, CC_box of 0.832, and Backbone F1 score of 0.728.

The third example (Figure 2g) is the metazoan retromer–sorting nexins 3 (SNX3) complex, modeled using a 9.5 Å cryo-EM map (EMD-12221; PDB ID: 7BLO^35^). This map did not resolve the C-terminal region of vacuolar protein sorting-associated protein 35 (VPS35). To fit the AF2 model of VPS35 to the map, only 320 out of 796 residues needed to be selected (right panel). DMcloud successfully modeled only the regions supported by the cryo-EM density, achieving the highest sequence coverage (0.876), CC_box (0.722), and backbone F1 score (0.878) among the methods tested (Extended Data Fig. 2c). In contrast, DiffModeler and EMBuild yielded substantially lower CC_box values (0.486 and 0.475, respectively) and backbone F1 scores (0.459 and 0.359, respectively), largely due to erroneous fitting of the unresolved C-terminal region of VPS35.

### Modeling performance on the high-resolution map dataset

Next, we evaluated DMcloud on a set of 50 high-resolution cryo-EM maps with resolutions ranging from 2.5 Å to 5.0 Å. This resolution range is most suitable for de novo modeling methods, as main-chain density is generally visible and traceable. Nevertheless, we tested DMcloud in this range to assess its applicability. In this dataset, the number of chains ranges from 1 to 35 (Supplementary Tables S4 and S5). Figure 3 summarizes the results and Supplementary Table S6 provides individual data. The average sequence coverage achieved by DMcloud (0.729) was on par with EMBuild (0.731), and both were substantially higher than DiffModeler (0.562), DEMO-EM2 (0.518), and ModelAngelo (0.558). In terms of CC_box and backbone F1 score, DMcloud outperformed EMBuild, with CC_box values of 0.559 versus 0.494 and backbone F1 scores of 0.737 versus 0.624, respectively.

**Figure 3.**
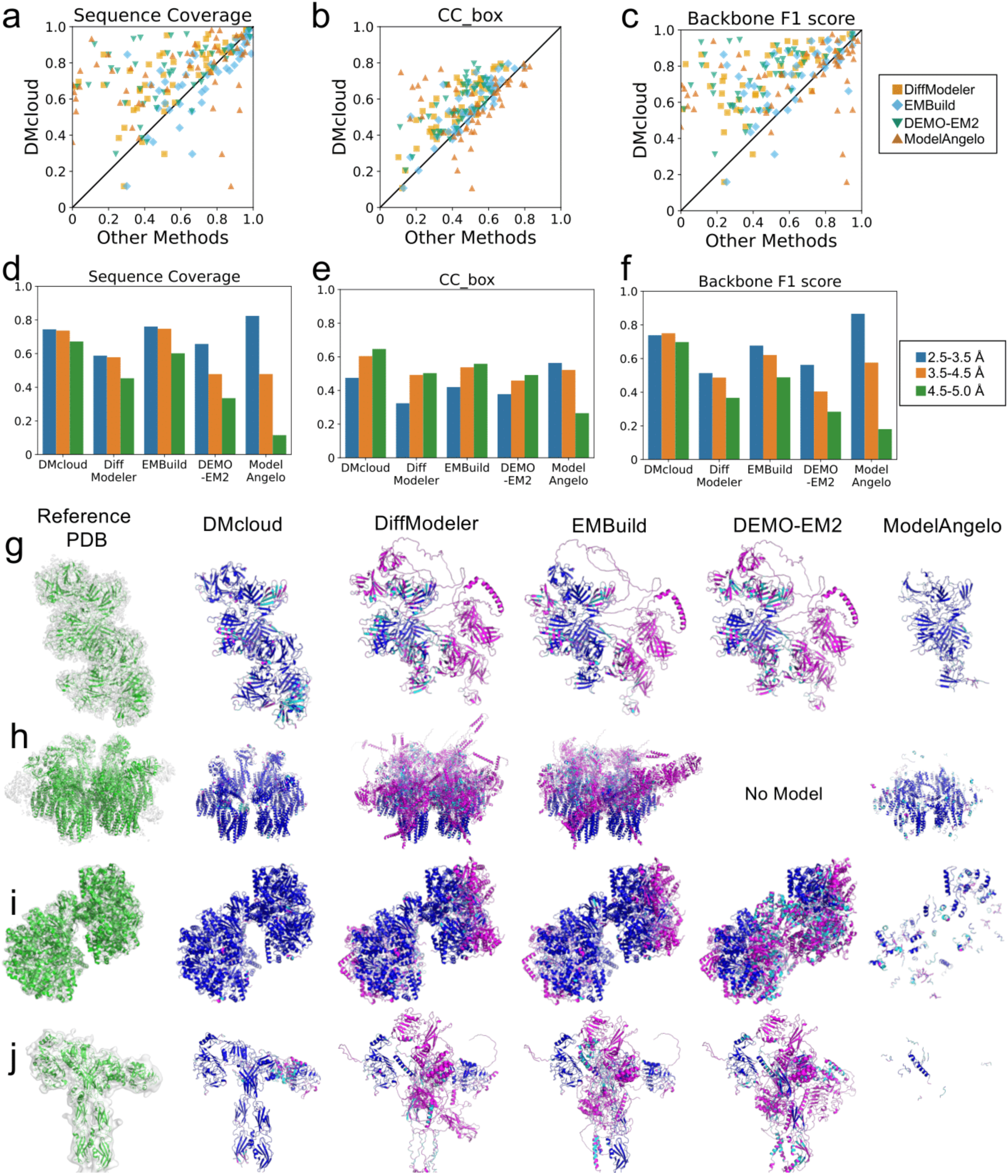
Modeling performance on 50 High-resolution cryo-EM map targets. a-c. Plots compare the performance of DMcloud (y-axis) with three other methods (x-axis), DiffModeler (orange squares), EMBuild (cyan diamonds), DEMO-EM2 (blue downward triangles), and ModelAngelo (brown upward triangles). **a.** Sequence Coverage, **b.** CC_box; **c.** F1 score. **d-f.** Breakdown of modeling performance across three resolution ranges: 2.5–3.5 Å (blue), 3.5–4.5 Å (orange), and 4.5–5.0 Å (green). **d.** Sequence Coverage, **e.** CC_box; **f.** F1 score. **g-i.** Model examples at different resolution ranges. Along with the PDB structure and the EM density map at the author-recommended contour level, models by the five methods are shown. Residues are colored using the same color scheme as in Figure 2. **g.** Insulin-like growth factor 2 receptor (EMD-20815; res. 3.46 Å, PDB 6UM1). single chain, 2,208 residues. Sequence Coverage 0.678, CC_box 0.484; Backbone F1 score 0.689. **h.** ESX-5 secretin system (EMD-12521; res. 4.48 Å; PDB 7NPU). 24 chains, 8,230 residues. Sequence Coverage 0.646, CC_box 0.512; Backbone F1 score 0.743. **i.** McrB-McrC complex without DNA binding domains (EMD-0315; res. 4.8 Å, PDB 6HZ9). 14 chains, 4,120 residues. Sequence Coverage 0.933, CC_box 0.766; Backbone F1 score 0.939. **j.** Mouse insulin receptor (EMD-28724; res. 4.9 Å, PDB 8EYY). 6 chains, 1,773 residues. Only the figure shows the extracellular region. Sequence Coverage 0.757, CC_box 0.718; Backbone F1 score 0.649.

Figures 3d to 3f show the performance of the methods broken down by resolution intervals of 0.5 Å. Extended Data Figure 3 and 4 have other evaluations including CC_mask, Sequence Match, Backbone precision, and Backbone recall. At higher resolutions (above 3.5 Å), DMcloud showed consistent performance across the three intervals even at the worst resolution range of 4.5 to 5.0 Å. In contrast, ModelAngelo exhibited a typical performance profile of de novo modeling methods, with high accuracy at the high-resolution range for which it was trained, followed by a sharp decline in performance at lower resolutions. The other three fitting methods exhibited a greater decline in sequence coverage and backbone F1 score than DMcloud as resolution decreased.

**Figure 4.**
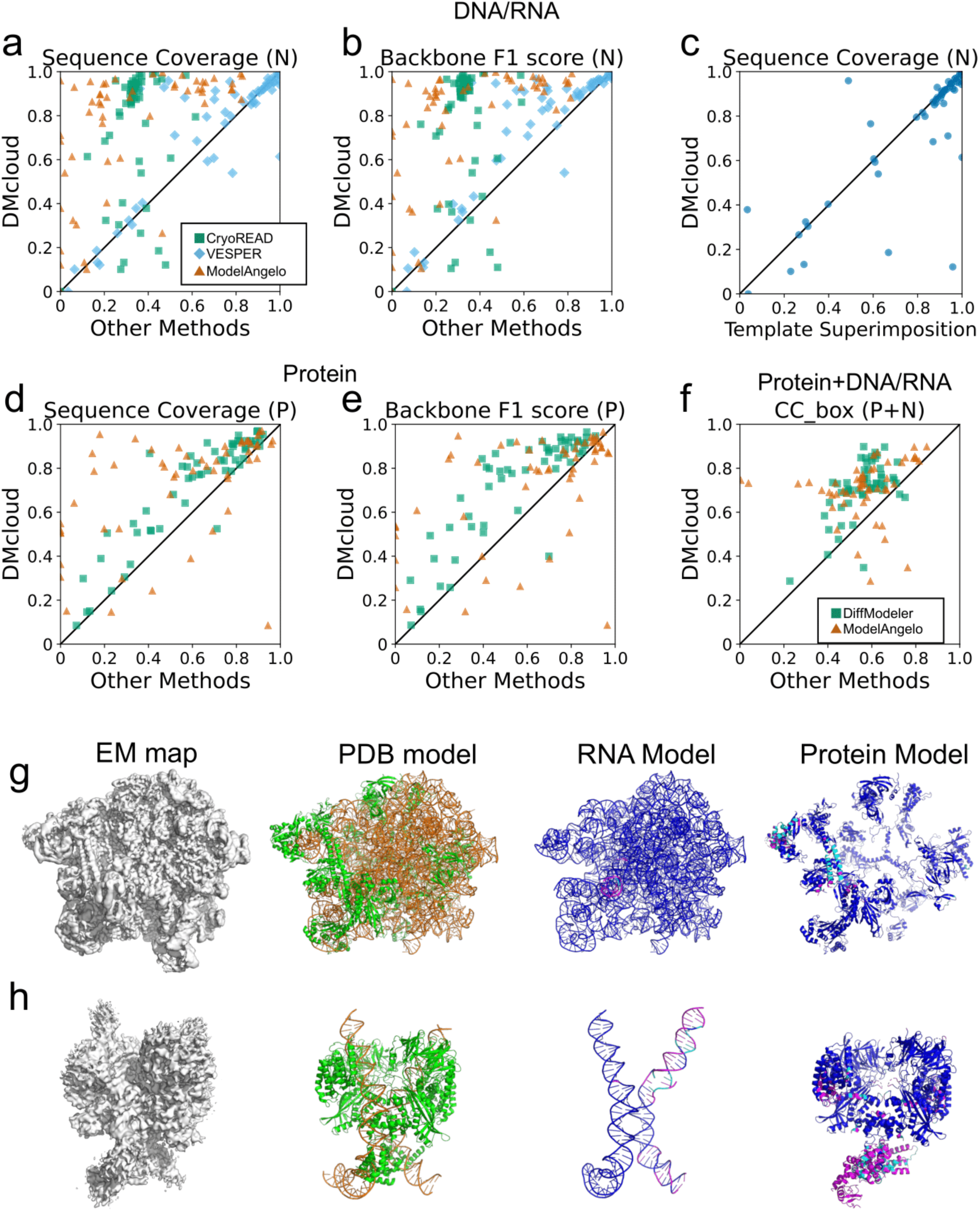
Modeling results on 55 maps with DNA/RNA-Protein complexes. a-f. Each plot compares DMcloud (y-axis) with other methods (x-axis). Consistent with the evaluation for protein modeling, a nucleotide in a model was considered as correct if it was placed within 3 Å to the correct position in the map. **a, b.** Comparison with CryoREAD, VESPER, and ModelAngelo for modeling DNA or RNA structures in terms of sequence coverage (SC) and the backbone F1 score. **c.** SC of DMcloud relative to the template model accuracy that was measured by superimposing the template structure to the PDB entry. with PyMol’s *align* command. **d, e**. Comparison with DiffModeler and ModelAngelo in protein modeling in terms of SC and backbone F1 score. **f.** CC_box of the DNA/RNA–protein complex models. **g.** Ribosomal 50S subunit with P-tRNA and RqcH (EMD-11889; res 2.9 Å, PDB: 7AS8). RNA: 2 chains; protein: 30 chains, 3,810 residues. Sequence coverage: 0.990 (RNA), 0.924 (protein). CC_box: 0.898 (RNA+protein). Backbone F1: 0.991 (RNA), 0.917 (protein). **h.** Mouse RAG1–RAG2 pre-reaction complex with DNA (EMD-20030; res. 3.6 Å, PDB: 6OEM). DNA: 4 chains, 206 residues; protein: 6 chains, 2,000 residues. Sequence coverage: 0.765 (DNA), 0.856 (protein). CC_box: 0.794 (DNA+protein). Backbone F1: 0.756 (DNA), 0.798 (protein). The models are colored using the same color scheme as in Figure 2, 3. Nucleotides (phosphates) are in blue if they are within 3 Å to their correct positions in the reference PDB; cyan indicates it is closer than 3 Å to a phosphate of a different nucleotide; magenta represents the predicted position deviates more than 3 Å from any phosphates in the reference PDB structure.

### Examples of high-resolution maps

Four illustrative examples from high-resolution maps are shown in Fig. 3g to 3j. The first example (Fig. 3g) is the insulin-like growth factor 2 receptor modeled with a cryo-EM map determined at a resolution of 3.46 Å (EMD-20815). This protein consists of 2,208 residues, which were segmented into 15 domains by the domain identification procedure in DMcloud (PDB ID: 6UM1^36^). The AF2 model of this protein shows large deviations in relative domain orientations, resulting in a TM-score of 0.585. Consequently, DiffModeler, which performs a global fit of AF2 models without local adjustments, was only able to place 382 residues within 3 Å of the reference (blue), leaving 17.3% of the regions outside of the density (magenta). EMBuild and DEMO-EM2, despite their ability to fit local domains, nonetheless produced models with similar inaccuracies to those of DiffModeler. At this high resolution, ModelAngelo was able to model only the core regions with locally high resolution, leaving the rest unmodeled. In contrast, DMcloud produced a more complete and accurate model. The sequence coverage and CC_box of the DMcloud model were 0.678 and 0.484, respectively. For comparison, the sequence coverage values for DiffModeler, EMBuild, DEMO-EM2, and ModelAngelo were 0.173, 0.433, 0.191, and 0.437, respectively, while their CC_box values were 0.199, 0.323, 0.225, and 0.510, respectively.

The next example (Fig. 3h) is from a map at a lower resolution than the previous one, 4.48 Å (EMD-12521, PDB ID: 7NPU^37^). This complex consists of 24 chains (two homo-hexamers and a homo-12 mer) with 8,230 residues. The AF2 models of the three different chains had varying accuracy: the model for chains C1-6 is accurate, with a TM-Score of 0.860, but those for chains B1-B6 and chains D1-DC were less accurate, with TM-scores of 0.657 and 0.728, respectively. DMcloud was able to identify and utilize the correct regions of the AF2 models, whereas DiffModeler and EMBuild incorrectly placed models containing inaccurate regions (highlighted in magenta). This difference is reflected in the backbone F1 scores: DMcloud achieved 0.743, while DiffModeler and EMBuild yielded substantially lower scores of 0.239 and 0.320, respectively. DEMO-EM2 failed to produce a structure, likely due to the large number of chains of this complex. The third example is the structure of McrB-McrC complex (PDB ID: 6HZ9^38^), modeled with a 4.8 Å resolution map (EMD-0315). This 14-chain complex consists of two different chains, McrB and McrC, 12 and two copies each. In this example, although the individual AF2 models are accurate (TM-scores of 0.965 to 0.987 and 0.927 for McrB and McrC, respectively), correctly placing the chains into the map was challenging due to their predominantly α-helical structures and the large number of copies. Nevertheless, DMcloud accurately reconstructed all chains without misplacement (sequence coverage: 0.933, CC_box: 0.766, backbone F1 score: 0.939). Several chains in the EMbuild and DEMO-EM2 models were misplaced (magenta), reducing the overall accuracy. At this resolution, models generated by ModelAngelo are highly fragmented.

The final example (Fig. 3j) is another structure model derived from a map at close to 5 Å resolution. (4.9 Å; EMD-28724). This structure of the mouse insulin receptor (PDB ID: 8EYY^39^) forms an asymmetric conformation containing six chains. The two largest chains (chain A and B) form a homodimer, but they have largely different conformations (Cα-RMSD: 15.1 Å). The AF2 model has different conformation from both chains, with a TM-score of 0.602 to chain A and 0.496 to chain B conformation. DMcloud achieved the best performance with sequence coverage of 0.757 by identifying and aligning consistent parts in the AF2 model to the map. In contrast, DiffModeler, EMBuild and DEMO-EM2 largely misplaced an AF2 model. ModelAngelo did not produce a meaningful structure.

These examples demonstrate that modeling remains challenging even for maps better than 5 Å resolution. Nevertheless, DMcloud consistently shows robustness and accuracy in modeling complex protein structures. Importantly, DMcloud relies solely on unique AF2 models, without prior knowledge of the number of copies of each chain in the map.

### Modeling DNA/RNA-Protein complexes

In this section, we demonstrate that DMcloud can fit nucleic acid structures when sufficiently accurate models are available. Unlike the protein modeling cases, where predicted structures were used for fitting, we performed sequence similarity searches using BLAST^40^ against the PDB to find nucleic acid structures from entries that were not identical to the target PDB entry. We did the structural search because currently there is no generally reliable nucleic acid structure prediction methods available. The dataset include 55 cryo-EM maps that were used in the test set of CryoREAD^32^ and DiffModeler^17^. The maps include 1 to 6 chains of DNAs or RNAs (57 to 4,286 nucleotides) with 1 to 87 protein structures (447 to 22,589 residues). Supplementary Table S7 and S8 provide the details of the dataset.

Figure 4a to 4e summarizes the modeling performance of DNA/RNA only (Fig. 4a, 4b, and 4c), protein only (Fig. 4d and 4e) and both (Fig. 4f) in the maps. Results of using additional metrics are shown in Extended Data Figure 5 and individual results are provided in Supplementary Table S9. As shown in Fig. 4a and 4b, DMcloud showed overall higher sequence recall and backbone F1 score for almost all the targets when compared to CryoREAD, VESPER, and ModelAngelo. The average sequence coverage (SC) and backbone F1 score of DMcloud were 0.754 and 0.768, respectively. The comparison with VESPER is particularly interesting, as both methods used the same nucleic acid structures for fitting. As shown, except for a few cases, DMcloud achieved higher SC values, indicating that its local fitting capability effectively produced more accurate models. In Fig. 4c, we evaluated the SC of nucleic acid models by DMcloud relative to the structure accuracy of the template structure to fit. Template sequence coverage on the x-axis was evaluated by superimposing template models onto the reference PDB and calculating SC. When template structures were highly accurate (SC > 0.8), DMcloud placed them in the correct pose in most cases (81.5%, 44 out of 54), achieving SC values within 0.05 of those obtained by direct superimposition. When the template structures are less accurate (SC < 0.8), there are a couple of cases where DMcloud was able to assemble locally correct parts of the templates, showing substantially higher SC than the superimposition. There were also opposite cases where the SC achieved by DMcloud was lower than the overall template accuracy. Such cases occurred when the template contained many locally similar structural regions that DMcloud misplaced (e.g., EMD-23530: template SC 0.958, DMcloud SC 0.121; EMD-24238: template SC 0.669, DMcloud SC 0.186), or when CryoREAD failed to correctly identify phosphate positions used for point cloud matching (e.g., EMD-8013: template SC 1.00, DMcloud SC 0.614). Proteins in the maps were modeled substantially more accurately by DMcloud than by two comparison methods (Fig. 4d, 4e), consistent with the results observed in the previous datasets (Fig. 2; Fig. 3). The modeling accuracy of the entire protein and nucleic acid complexes were shown in Fig. 4f. Among the two methods shown in the plot, DMcloud clearly showed the highest sequence coverage for almost all the target maps.

### Modeling examples of DNA/RNA-protein complexes

Figures 4g and 4h show two illustrative examples in the DNA/RNA-protein dataset. The first example is a complex of the Ribosomal 50S subunit with P-tRNA and RqcH (Figure 4f). The cryo-EM map EMD-11889 (2.9 Å resolution) with PDB ID 7AS8^41^ includes two RNA components (50S rRNA and tRNA) and 30 chains of proteins. DMcloud achieved high accuracy (sequence coverage, SC 0.990; backbone F1 score 0.991). VESPER also achieved similar performance (SC 0.987, backbone F1 score 0.987). The other two methods, CryoREAD, and ModelAngelo, showed lower RNA modeling accuracy (SC 0.359 and 0.821; backbone F1 score 0.325 and 0.818, respectively). In contrast, for protein modeling, DMcloud clearly performed best (SC 0.924; F1 0.917), surpassing DiffModeler (SC 0.807; F1 0.788) and ModelAngelo (SC 0.824; F1 0.895). At the RNA-protein complex level, DMcloud also gave the highest model-map agreement (CC_box 0.898, CC_mask 0.851) relative to DiffModeler (0.563, 0.401) and ModelAngelo (0.850, 0.755).

The second example is the mouse RAG1–RAG2 pre-reaction complex with DNA (Fig. 4h; EMD-20030, res. 3.6 Å; PDB 6OEM^42^), a macromolecule that consists of 4 DNA chains (206 nucleotides) and 6 chains of protein (2,000 residues). On this target, DMcloud achieved the best DNA modeling accuracy (sequence coverage, SC 0.765; backbone F1 0.756), whereas VESPER, CryoREAD, and ModelAngelo were lower (SC 0.750, 0.388, 0.191; F1 0.750, 0.388, 0.227, respectively). DMcloud and VESPER used the same template structures but DMcloud’s point-cloud fitting approach enabled a more precise alignment. For protein modeling, DMcloud again performed best (SC 0.856) outperforming DiffModeler, which was the second best (SC 0.592), and the other methods.

### Large Complex Targets

In this section, we discuss DMcloud’s models for large assemblies with many chains and diverse stoichiometries (Fig. 5). The first example (Fig. 5a) is the mitochondrial heat shock protein 60 (mtHsp60)-mitochondrial heat shock protein 10 (mtHsp10) focus structure (EMD-29818, res. 3.4 Å; PDB: 8G7O^43^), included in the high-resolution dataset (Fig. 3). The authors constructed six maps in their work^43^. This PDB model includes only the upper half of the complex, consisting of seven copies each of mtHsp10 and mtHsp60, because the corresponding map (EMD-29818) was obtained through focused refinement of the upper region, leaving the lower part at lower resolution. Remarkably, DMcloud reconstructed not only the upper region but also accurately assembled the lower half of the complex as shown in the figure. The upper region of the DMcloud model achieved a sequence coverage (SC) of 0.958, while the full model, including the lower part, reached an SC of 0.884 compared with the complete mtHsp60–mtHsp10 complex (PDB 8G7N), which was built from a different map (EMD-29817). Notably, DMcloud automatically detected the symmetric subunit arrangement using only two unique AF2 models and without any prior information on stoichiometry, despite the large size of the complex.

**Figure 5.**
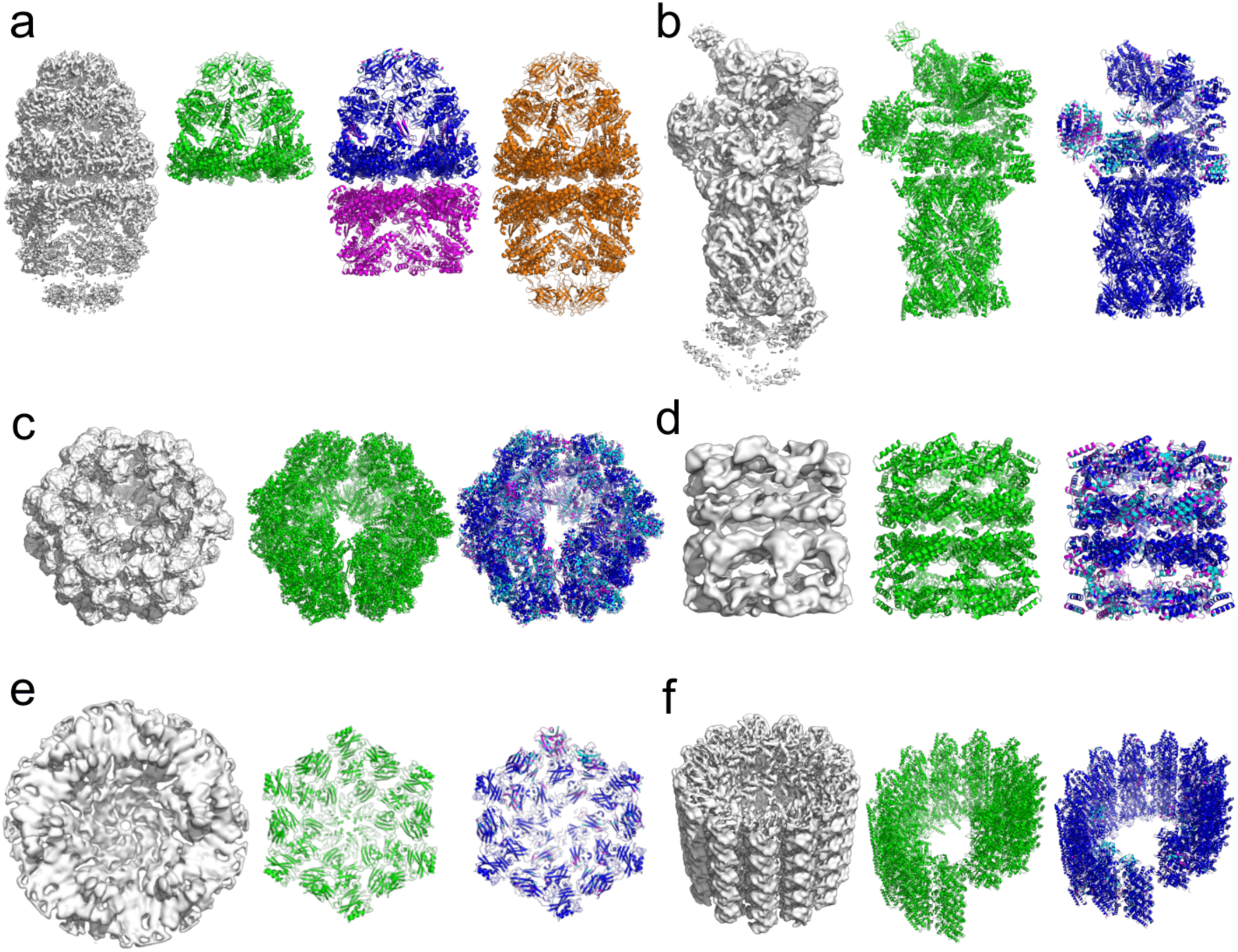
Examples of modeling for large macromolecular assemblies. For each example, three figures are shown: the cryo-EM density map at the recommended contour level; the reference PDB structure model; and the model constructed by DMcloud. DMcloud models are colored in blue, cyan, and magenta depending on the overlap with the PDB model using the same color scheme as Figs. 2-4. **a.** mtHsp60-mtHsp10 focus structure (EMD-29818, res. 3.4 Å; PDB: 8G7O). 14 chains, 4,389 residues. Sequence Coverage (SC) 0.958, CC_box 0.681; Backbone F1 score 0.662. The structure model on the right in orange shows the mtHsp60-mtHsp10 full complex structure (PDB 8G7N) built from a different map (EMD-29817). The bottom half of the DMcloud model is colored in magenta because it was compared with 8G7O, which lacks this part. See the text. **b.** 26S proteasome (EMD-4321, res. 5.4 Å, PDB: 6FVV). 47 chains, 14,015 residues. SC 0.732, CC_box 0.643, Backbone F1 score 0.764. c. *Lumbricus terrestris* hemoglobin (EMD-2627, res. 8.1 Å; PDB 4V93). 180 chains, 11,664 residues. SC 0.481, CC_box 0.695, and backbone F1 score of 0.495. **d.** Apo GroEL (EMD-1042, res. 10.3 Å, PDB 1GR5). 14 chains, 7,238 residues. SC 0.526, CC_box 0.827, Backbone F1 score 0.537. For these two cryo-ET targets in panels **e** and **f**, protein regions in the maps were segmented before using DMcloud. **e.** S-layer of *Haloferax volcanii* (EMD-13637, res. 8.0 Å, PDB 7PTT, cryo-ET map). 6 chains, 4,350 residues. SC 0.683, CC_box 0.658, Backbone F1 score 0.697. **f.** A microtubule complex with intraluminal proteins (EMD-26019, cryo-ET map, res. 6.7 Å; PDB 7TNS) 101 chains, 27,593 residues. SC 0.761, CC_box 0.349, Backbone F1 score 0.820.

The second example (Fig. 5b) is 26S proteasome (EMD-4321, res. 5.4 Å; PDB 6FVV^44^), which consists of 47 chains and 14,015 residues. We present this example because the proteasome complex of 47 chains was previously highlighted in the DiffModeler paper^17^ as a successful case of modeling a large complex. DMcloud produced a more accurate model with an SC of 0.732, CC_box of 0.643, and backbone F1 score of 0.764 in comparison to the DiffModeler model (SC: 0.641; CC_box: 0.545; backbone F1 score: 0.616).

The third example (Fig. 5c) is a giant 3.5 MDa hexagonal-bilayer complex, *Lumbricus terrestris* hemoglobin (EMD-2627, res. 8.1 Å; PDB 4V93^45^). It consists of 180 chains, including 144 globin subunits with four types and 36 linker chains. This is a challenging target because the four hemoglobin chains have almost identical structures (the average RMSD of the four types of hemoglobin is 1.60 Å) yet they share less than 46.4% sequence identity. Because the method primarily relies on backbone shape for structure fitting, it can misassign subunits when they share highly similar folds but have low sequence similarity. The DMcloud model achieved a CC_box of 0.695; however, its sequence correspondence dropped to 0.481. DiffModeler performed worse, with an SC of 0.232, while the other two methods (EMBuild and DEMO-EM2) performed even worse.

The next one is an example of a low-resolution map of 10.3 Å (Fig. 5d). It is apo GroEL (EMD-1042, PDB 1GR5^46^), which consists of a homo-14-mer complex with 7,238 residues. At resolutions lower than 10 Å, cryo-EM maps usually reveal only coarse structural features such as domain shapes and helical bundles, making model fitting very challenging. Remarkably, DMcloud correctly identified the symmetric arrangement of all 14 subunits using a single unique AF2 model, reflecting a high CC_box value of 0.827.

Additionally, we applied DMcloud on two cryo-ET maps. Fig. 5d is hexameric S-layer of *Haloferax volcanii* modelled from a cryo-ET map (EMD-13637, res. 8.0 Å; PDB 7PTT^47^), which consists of homo hexamer with 4,350 residues. Each chain has six repeated immunoglobulin-like domains^47^. DMcloud was able to place the AF2 model of the chain in the correct, symmetrical positions despite the low resolution of the map. CC_box was 0.658, and SC was 0.683.

The last panel is a cryo-ET map of a microtubule complex with intraluminal proteins (EMD-26019, res. 6.7 Å; PDB 7TNS^48^). This large consists of 101 chains and 27,593 residues. Similar to Fig. 5c, a key challenge in structure fitting for this map is that the complex consists of two structurally very similar subunits, α- and β-tubulins (TM-score: 0.92), which share only 42% sequence identity. The complex is organized into alternating layers of α- and β-tubulins. It turned out that DMcloud fitted α-tubulin structure to most of the places, which resulted in a backbone F1 score of 0.621 and SC of 0.576. But the modeling result drastically improved by specifying α- and β-tubulin layers, with backbone F1 of 0.820 and SC of 0.761. The examples in Fig. 5c and 5f illustrate a limitation of DMcloud: since it fits chains based primarily on main-chain conformation, subunits with nearly identical structures but different sequences cannot be well distinguished.

## Discussion

We present DMcloud, a new structure modeling approach for cryo-EM maps with resolutions up to approximately 10 Å. DMcloud performs accurate local fitting of protein and nucleic acid models to experimental maps by employing a diffusion model that emphasizes main-chain positions, combined with a series of point-cloud matching and clustering algorithms. By converting both cryo-EM maps and molecular models into point clouds, DMcloud selectively identifies local regions in the models that are consistent with the map density. This approach provides a major advantage over existing fitting methods, as it can effectively exclude incorrectly modeled regions from the fitting process. Despite the rapid advances in structure prediction, it remains common for predicted models to contain domains with incorrect orientations or long loops that fail to fit the target cryo-EM density. DMcloud can identify and remove such conformationally inconsistent regions, leading to more accurate and interpretable model fitting. DMcloud consistently outperformed existing methods across datasets spanning medium to high resolution cryo-EM maps. Moreover, the method scales efficiently to large complexes containing numerous subunits. Unlike other similar methods, DMcloud does not require users to specify the number of copies for each chain. Instead, users only need to provide unique structure models, and DMcloud automatically identifies the number of copies by fitting each structure to the map.

Although DMcloud offers strong advantages over existing methods, it also has certain algorithmic limitations. First, as DMcloud is primarily a structure fitting method, it does not perform fine adjustments of atomic positions within the fitted models, making it difficult to fully optimize the fit to the map. In addition, because DMcloud relies on backbone positions identified by the diffusion model, fitting becomes unreliable when the backbone cannot be accurately detected, such as in local regions in maps with particularly low resolution.

These limitations could be addressed by integrating fully flexible point-to-point registration, incorporating a local structural refinement step guided by the cryo-EM density, and improving the accuracy of backbone position detection. In particular, the local refinement process could adjust backbone and side-chain conformations to enhance model accuracy, especially in regions requiring small conformational corrections. Furthermore, similar to how structure models predicted by AlphaFold2 are utilized in DeepMainmast^6^, structure prediction could be more directly integrated into the modeling pipeline to further improve fitting performance. Extending the methodology to cryo-electron tomography (cryo-ET) data also remains an important direction for future work.

## Methods

### Diffusion Model for cryo-EM maps

To enhance the cryo-EM map for accurate structure fitting, DMcloud modifies the cryo-EM map using the diffusion model that was developed for DiffModeler^17^. The cryo-EM maps is subjected to preprocessing before being used in the diffusion model. If a map has a grid size different from 1.0 Å, it is interpolated to a 1.0 Å grid using trilinear interpolation. The density values within a map are normalized to the range of 0.0 to 1.0 by applying minimum-maximum normalization. Negative values are set to 0, and 0 is used as the minimum value for normalization. The maximum value for normalization is set to the 98th percentile density value, and any values above this are capped at 1.0. Boxes of 64³ Å³ are then extracted by scanning the map with a stride of 32 Å. The goal of the diffusion model is to generate backbone labels at positions of N, Cα, and C of protein backbone in the cryo-EM map. The denoising diffusion implicit model (DDIM) was used. It was trained on 230 maps and validated on 36 maps that have a resolution between 5 to 10 Å. For more details refer to the DiffModeler^17^ paper. Two examples of enhanced maps computed by the diffusion model are shown in the Extended Data Figure 6.

### Converting a diffusion model-processed map into point cloud of LDPs

The map processed by the diffusion model, where backbone features are enhanced, is represented as a point cloud composed of local dense points (LDPs), which serve as representative markers of high-density regions. As introduced in our earlier work^23^, a mean-shift algorithm is used to identify these LDPs from the enhanced density map. First, all grid points with a density value larger than 0.01 were selected in the map. The coordinates of each identified grid point *v* are iteratively updated using the neighboring grid points *N*(*v*) as:

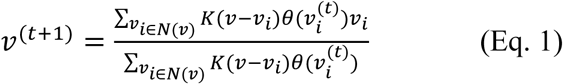

where 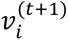 is the updated grid position of *v_i_*, K() is a Gaussian kernel function, *θ*(*v*) is the density value of the grid point *v*. The neighboring grid points *N*(*v*) is a set of grid points that satisfy ‖*v_i_* − *v*‖, ≤ 2 ∗ *σ*, *σ* is a bandwidth set to 2.0. The Gaussian kernel function K() is defined as:

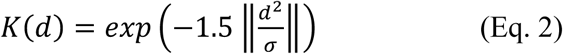

*d* is the Euclidean distance between the points, and *σ* is a bandwidth parameter. The mean shifting using Eq. 1 is iterated until convergence when ‖*v*^(*t*+1)^ − *v*^(*t*)^‖_2_ ≤ 0.001. After the mean-shifting process, points within 2.0 Å of each other are clustered together. From each cluster, the grid point with the highest density is selected as an LDP.

### Processing input protein structure models

Input protein structure models are also transformed into point cloud of positions of Cα, N, and C atoms of the main chain. It is performed in two steps: identification of domain structures, and extraction of points from the domains. For domain identification, we used DBSCAN. DBSCAN is a clustering algorithm that groups points based on their weights and special distribution. DBSCAN uses two parameters, *ε* and *min_total_weights*. *ε* represents the maximum distance between two points for them to be considered as neighbors. It is set to 5 Å. *min_total_weights* is the minimum total weights among neighbors for a point to be considered as a core point, which is set to 500. We used pLDDT confidence score from AF2 as the weight of each atom. Atoms that belong to residues with a pLDDT value below 50.0 are discarded.

Atoms (points) are classified into three categories: core, border, and noise. A point is classified as a *core point* if it has at least min_total_weights total weight among its neighbors within an ε Å radius. If a point has a lower total weight than required for a core point but lies within ε Å of a core point, it is classified as a *border point*. Points that meet neither criterion are labeled as *noise* and are discarded. Once classification is complete, DBSCAN initiates clustering using core points. Starting from a core point, the algorithm expands the cluster by recursively adding all neighboring core points within a 5 Å radius. Subsequently, border points that are neighbors of core points in the cluster are also included. This process continues until no additional points can be added. The algorithm then proceeds to a new unvisited core point, repeating the process until all points have been assigned to clusters or labeled as noise.

### Point cloud matching between protein domains and the processed map

Similar to the identification of domains in protein models performed in the previous step, the diffusion-model processed map is also segmented into local regions. Then, each point cloud that represents individual domain structure is matched to individual regions of the map. LDPs in the map are segmented into local groups as follows in two steps: First, representative LDPs are selected from the entire LDPs of the map by point down-sampling with a 10 Å interval. Then, LDPs within a 15 Å distance for each representative points are grouped and assigned to the representative point. Then, each segmented LDPs around a representative one is matched to each of protein domains using cloud matching with RANSAC followed by ICP.

RANSAC aligns two point clouds by considering similarity of features of points in the clouds. We characterized points in the clouds with Fast Point Feature Histograms^29^ (FPFH), which were computed using the open3d package^49^. FPFH is a feature descriptor that represents the local geometric properties of a point in a 33-dimensional space (See the subsequent section). For each point in the domain structure, five most similar points in the segmented LDPs of the map based on the FPFH similarity are recorded as a list of pairs. Then, three random pairs of points are selected from the list iteratively. For each iteration, rotation and translation matrices are computed to align the selected three point pairs. Then, this transformation is applied to the entire points in the domain structure, and points where the distance to their corresponding LDPs within 3 Å are considered as an initial alignment. This initial alignment is refined using ICP, which aligns the entire set of LDPs of the whole map with the point cloud of the protein domain structure. ICP is an iterative optimization method that minimizes the difference between two point clouds. Starting from the matched point pairs in the initial alignment, the optimal rotation and translation were computed that minimize the mean square error between them. After applying the transformation, new corresponding point pairs that are within a cutoff distance of *r* Å are added to the alignment. At the same time, point pairs that become more than *r* Å apart are removed from the alignment. For protein structures, we set *r* = 3.0 Å, and for DNA/RNA structures, r = 4.0 Å. ICP repeats this iteration until convergence. Finally, DBSCAN is applied to the alignment to identify clusters and remove noise. The size and density of the clusters are controlled by its two parameters, epsilon (*ε*), which defines the radius of the local neighbors around each point, and *min_total_weights*, which defines the minimum total weight for a core point. To construct clusters of various sizes and densities, DMcloud calculates clusters using exhaustive combinations of *ε* = [3, 4, 5] and *total_min_weights* = [10, 20, 30]. The *total_min_weights* parameter considers the LDDT score for each aligned point (see the later section). Among the clusters identified by DBSCAN, clusters were removed if they have an average LDDDT (aveLDDT) of less than 0.4 or have fewer than 200 aligned points (100 for DNA/RNA-protein complex targets), as they were considered as of a low quality. Remaining clusters of alignments are recorded as candidates of fitted models in the structure pool.

### Fast Point Feature Histogram (FPFH)

FPFH is a descriptor that captures local geometric features of points in a point cloud^29^. FPFH describes distributions of three types of angles computed for the point of interest and neighboring points around it (Extended Data Figure 7). FPFH is a weighted sum of a histogram feature, named as Simplified Point Feature Histogram (SPFH), computed for each point. SPFH calculates three key angular features between a query point and its neighbors within a set radius. Three angles are: alpha (α), the angle between the line connecting the query point to a neighbor point and the query point’s surface normal; phi (φ), the angle between the direction vector from the query point to the neighbor point and the neighbor point’s surface normal; and theta (θ), which measures the angular difference between the normals of the query point and the neighbor point. These features are divided into 11 bins each, creating a 33-dimensional histogram that represents the local geometry around the point. For a protein structure, we used the radius of 8 Å, while radii of 8 Å, 14 Å, and 20 Å were used for DNA/RNA structures. Using SPFH computed for each point, FPFH combines SPFH of *k* neighboring points (*p_k_*) within the radius cutoff with weights defined as the inverse of the distance between the query point (*p*) and the neighbors:

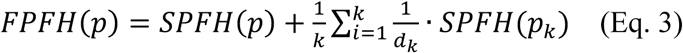

where d_k_ is the distance between *p* and p_k_. For DNA/RNA targets, FPFH are computed using three radii (8, 14, and 20 Å). Based on these features, three corresponding structure models are generated. Among the three models, the model with the highest rawLDDT score is selected as the final prediction.

### LDDT score used for evaluating protein domain fit to a map

The LDDT (Local Distance Difference Test) score^31^ was originally designed to compare local structure environment of a residue in a computational protein model with the one in the native protein structure. Here, we use LDDT to evaluate domain structure fit to LDPs in a map.

For a given point *i* in the domain structure model, its neighboring points, *j* ∈ *G_i_*, within a cutoff distance *R*_cutoff_ (= 8 Å for protein structures, 8, 14, 20 Å for DNA/RNA structures) in the domain structure, are identified. The corresponding aligned point in LDPs for *i* is denoted as *I* while the corresponding point for *j* is denoted as *J*. For each pair of *i* and *j* ∈ *G*_i_, the distance *d_ij_^model^* between *i* and *j*, and corresponding distance *d_I,J_^LDPs^* between *I* and *J*, were computed, and the difference between these distances, *Δ_ij_* is compute as Δd_ij_=∣d_i,j_^model^−d_I,J_^LDPs^∣. Using a set of distance difference thresholds [0.5, 1.0, 2.0, 4.0], the score of pair *S_i,j_* is computed as

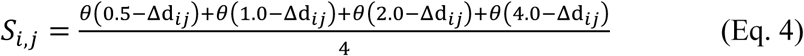

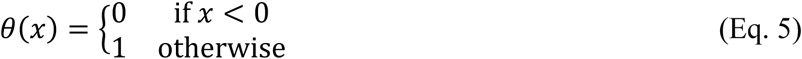

Then, the LDDT score of point *i* with the corresponding point *I* is computed as:

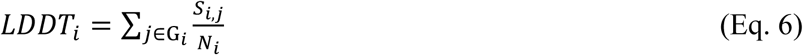

where *N_i_* is the number of neighboring points of *i*. The LDDT score ranges from 0 to 1.0 with 1.0 showing a perfect agreement of the point *i* and *I*. Using *LDDT_i_,* the fitness and coverage between the fitted domain structure and LDPs are computed. We defined rawLDDT as the sum of :

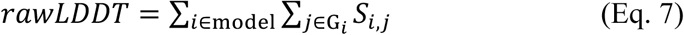

where *i* is all points in the fitted domain structure. Thus, *rawLDDT* is the sum of *LDDT_i_* of the structure domain. We also define the average of *rawLDDT* score (*aveLDDT*) as

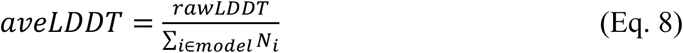

The definition of *aveLDDT* differs from the global LDDT score used in AlphaFold. In the global LDDT, the score is computed by averaging the residue-wise *LDDT_i_* values. In *aveLDDT*, we first sum up the pairwise agreement scores *S_i,j_*, and then normalize by the total number of residue pairs in the model.

### Assembling fitted models using CP Solver

Fitted structure domains are selected and combined using a CP solver. The CP solver optimizes a combination of domains under a constraint. Here, the constraint is to maximize the sum of the domain structure fitness and the penalty if selected domains overlap:

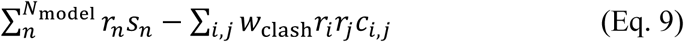

In the first term, *Nmodel* is the number of domain structure models fitted to the map and *s_n_* (*n* = 1, …, *N*_model_) is the rawLDDT score of each fitted model. *r*_*n*_∈ {0, 1} indicates whether the domain structure *n* is included in the assembly. In the second term, *c_i,j_* is the number of clashes between two models, *i* and *j.* A clash is defined between two domains when aligned LDPs with the two domains are within 2.0 Å to each other. *w*_clash_ is a weight of penalties, which is set to 30. If two domains *i* and *j* have more than 50% of their LDPs clashing, then the combination is considered invalid and cannot include both domains, i.e.

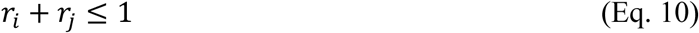

### Copying full atoms to aligned and neighboring residues

The assembled models so far include only backbone atoms (N, Cα, C; oxygen not included) of aligned domains. This final step will copy all the atoms in residues from the structure model if the residues meet one of two following conditions: (1) at least one backbone atom in the residue aligns with an LDP; or (2) at least one backbone atom of the residue is located within 5 Å of other aligned backbone atoms.

### DNA/RNA structure fitting

DNA/RNA structure models taken from PDB are fit to predicted sugar and phosphate positions in the target cryo-EM map. Prediction of sugar positions, concretely, five key sugar atoms (C1’, C2’, C3’, C4’, and O4) and phosphate atoms, is performed using a deep learning model used in CryoREAD. Accordingly, DNA and RNA structures are represented by two-color (i.e. sugar and phosphate) point clouds. Similar to the protein structure model fitting, the generated two-color point clouds are segmented into domains using DBSCAN with parameters set to ε = 15 Å and *min_total_weights* = 5. The weight of each point is initialized to 1.0.

### Two-color Point Cloud Matching

In the two-color point cloud matching step, the matching process is customized to differentiate between sugar and phosphate points. During the RANSAC process, the five most similar points are selected based on FPFH similarity, considering the type of points, either sugar or phosphate. This approach ensures that only points of the same type (sugar to sugar, phosphate to phosphate) are paired during matching. Subsequent refinement using ICP also considers the type-specific alignment, optimizing the transformation to align sugar points in the model exclusively with sugar points in the LDPs, and phosphate points with phosphate points. Sugar or phosphate points in a DNA/RNA model that are within 4 Å to corresponding LDPs are considered aligned. To capture a diverse range of local geometric features, we compute the FPFH using three different local radii, 8, 14, and 20 Å. Among all the fitted models generated including ones with FPFH of different radii, the model with the highest total score is selected as the final output. The total score is defined as the *rawLDDT* score, which is computed from the single alignment in which sugar points in the model are aligned to sugar points in the LDPs, and phosphate points are aligned to phosphate points. In this computation, the score of pairs (*S_i,j_*) is calculated over all point pairs in the alignment. Therefore, sugar-sugar, sugar-phosphate, and sugar-phosphate distances contribute to *S_i,j_*. The rawLDDT indicates the overall geometric consistency between the model and the LDPs of sugar and phosphate.

### Constructing the dataset of high-resolution targets

We tested DMcloud on three datasets, as shown in Figs. 2, 3, and 4, which correspond to intermediate-resolution maps, high-resolution maps, and nucleic acid maps, respectively. The intermediate-resolution map dataset was taken from the DiffModeler paper^17^. The high-resolution dataset consisted of 50 cryo-EM maps collected from the Electron Microscopy Data Bank (EMDB) as of September 30, 2024, according to the following criteria: each map had a resolution between 2.5 Å and 5.0 Å and a corresponding deposited structure in the Protein Data Bank (PDB)^50,51^, containing at least 20 residues. Maps were excluded if the associated PDB structures had a cross-correlation coefficient or overlap score below 0.65 compared with the EM map. To eliminate redundancy with the *DiffModeler* training dataset, we computed pairwise global sequence identities between chains. Any map containing a chain with ≥25% sequence identity to a chain in the *DiffModeler* training set was discarded. From the remaining 509 maps, we randomly selected 10 models from each of the following resolution ranges: 2.5–3.0 Å, 3.0–3.5 Å, 3.5–4.0 Å, 4.0–4.5 Å, and 4.5–5.0 Å, resulting in a final curated set of 50 high-resolution cryo-EM maps.

### Constructing the dataset of DNA/RNA-protein targets

The 61 maps with DNA/RNA-protein complex were originally used as the test dataset in the CryoREAD paper^32^ and the DiffModeler paper^17^. The dataset was constructed through the following process: First, cryo-EM maps were selected from the EMDB on June 10, 2021, with resolutions between 2.0 Å and 5.0 Å, each having an associated PDB entry with nucleic acid structures. The selection criteria required a minimum cross-correlation of 0.65 between the density of the cryo-EM map and the corresponding PDB-derived simulated map to ensure alignment quality. To reduce redundancy, maps were clustered based on sequence identity, with clustering criteria of at least 30% sequence identity for protein chains and 80% for DNA/RNA chains. From these clusters, 68 targets were randomly selected. Chains shorter than 50 nucleotides or containing any unknown residues (X) were excluded from the dataset.

To identify known template structures for each target, we constructed a DNA/RNA sequence database from PDB entries as of February 17, 2024, comprising 865,773 DNA/RNA chains. For six targets (EMD-6441, EMD-4138, EMD-12973, EMD-20452, EMD-3817, and EMD-30005), the BLAST search did not identify template structures using an e-value cutoff of 10.0. We removed these 5 targets from the benchmark dataset. For protein modeling, we used the same AlphaFold Database models used in the DiffModeler paper^17^. The final benchmark dataset consisted of 55 DNA/RNA–protein complex maps. For one target (EMD-12177), DMcloud could not generate a model due to a lack of structural similarity between the template and the cryo-EM density. For this target, only the protein model was evaluated.

### Computational Time

The computational time of DMcloud is reported in Extended Data Figure 8. We used a single GPU card (Nvidia RTX A6000 with 48 GB memory) and a CPU (Intel Xeon E5-1650 with 12 cores). In panels a and b, blue plots represent the computational time for backbone position detection by the diffusion model in the maps, while orange plots indicate the total computational time of DMcloud, including backbone position detection, point-cloud matching, and model assembly. In panel c, blue plots show the computational time for phosphate and sugar position detection in the maps using the neural network (3D-CNN) in the CryoREAD modeling pipeline, while orange plots represent the total computational time of DMcloud for DNA/RNA modeling. The total computational time depends on the number of detected points (backbone atoms, phosphate, and sugar), the number of chains, and the number of residues. For protein targets, the average computational time for the maps with 5,000-15,000 residues is 18.7 hours. For DNA/RNA maps of 2,500-3500 nucleotides, the average total computational time is 4.5 hours. The detailed data are provided in Supplementary Table S11.

### Evaluation Metrics

The constructed models by DMcloud and the other methods were evaluated by comparing them to the deposited PDB structure using the *phenix.chain_comparison* tool in Phenix^33^. Sequence coverage is defined as the fraction of Cα atoms in the deposited PDB structure that are correctly matched to the corresponding amino acid type in the predicted model within 3 Å. Sequence match represents the fraction of correctly predicted Cα atom positions and amino acid types among the correctly predicted Cα atom positions. Backbone recall is the fraction of Cα atoms in the deposited PDB structure that are correctly predicted within 3 Å, regardless of amino acid type. Backbone precision is the fraction of Cα atoms in the predicted model that are correctly predicted within 3 Å, also independent of amino acid type. For DNA/RNA models, phosphate atom positions are used for the evaluation instead of Cα atoms with 3 Å distance cutoff.

Using the backbone recall and precision, we also computed backbone F1 score, which combines the two values:

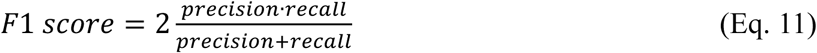

We computed two types of cross-correlation values between a structure model and the map, CC_mask and CC_box, using *phenix.map_model_cc* in Phenix. CC_mask is the correlation coefficient between the model-derived map and the experimental map inside the mask region built around the model, while CC_box is the correlation coefficient between the model-derived map and the whole experimental map.

### Structure models by the other methods DiffModeler

The predicted models for the Intermediate resolution dataset (71 maps) were obtained from https://zenodo.org/records/12155184. For the high-resolution dataset (50 maps) and DNA/RNA-protein complex targets (55 maps), we used the code v2.2, downloaded from https://github.com/kiharalab/DiffModeler. For targets that contain both nucleic acids and proteins, we first constructed the DNA/RNA models using CryoREAD and then performed DiffModeler for the protein components^17^.

### DEMO-EM2

We used the DEMO-EM2 code downloaded from https://zhanggroup.org/DEMO-EM/DEMO-EM2/. During the calculations for DEMO-EM2, we set the -rsc option to 0.1, because the computation with the default parameter took more than 72 hours for a single target for most of the targets.

### EMBuild

We used the EMbuild v1.1 code downloaded from http://huanglab.phys.hust.edu.cn/EMBuild/EMBuild_v1.1.tgz.

### ModelAngelo

We used the code version 1.0.12, downloaded from https://github.com/3dem/model-angelo

### VESPER

We used a GPU-version of VESPER. The code was downloaded from https://github.com/kiharalab/DiffModeler/tree/main/VESPER_CUDA

### CryoREAD

We used the code version 11.0, downloaded from https://github.com/kiharalab/CryoREAD.

### Captions of Supplementary Tables

Supplementary tables are provided in a separate Excel file.

**Supplementary Table S1. The benchmark dataset of Intermediate resolution maps.** The first column, “EMD-ID”, lists the EMDB IDs of the cryo-EM maps used in the benchmark dataset. The second column, ’PDB-ID,’ provides the PDB IDs of the structures associated with each map. The third column, “Resolution (Å)”, indicates the reported resolution of the deposited cryo-EM map. The fourth column, “Nchain”, specifies the number of chains in the corresponding PDB structure. The fifth column, ’Nresidues,’ shows the total number of residues in the PDB structure.

**Supplementary Table S2. AlphaFold2 single-chain structures for the intermediate resolution datasets.** The “EMD-ID” and “PDB-ID” columns represent the EMDB and PDB identifiers for each entry, respectively. The Resolution column lists the reported resolution of the deposited cryo-EM map. The “Chain_ID” column specifies the chain identifiers within the corresponding PDB structure, while the “Uniprot_ID “column provides the UniProt ID linked to the single-chain protein entry in the AlphaFold database. For cases marked as “AFDB search”, we reported the entry ID from the AlphaFold Database. The “TM-Score” column reports the TM-Score metric, which measures the structural similarity between the AlphaFold2 model and the native structure.

**Supplementary Table S3. Modeling performance on the intermediate resolution dataset.** Each row represents a specific cryo-EM map identified by its EMD-ID. We reported seven metrics (Sequence Coverage, CC_box, F1 Score, CC_mask, Sequence Match, Backbone Precision, and Backbone Recall). We analyzed five methods, DMcloud, DiffModeler, EMBuild, DEMO-EM2, and ModelAngelo across multiple evaluation metrics. Each method’s performance is reported under the respective column for each metric. For reference, “AF2 Model Superimposition” represents models obtained by directly superimposing AlphaFold Database models onto the reference PDB structures. The blank data corresponds to the method that failed to build the model for the target.

**Supplementary Table S4. Benchmark dataset of the high-resolution maps** The first column, “EMD-ID”, lists the EMDB IDs of the cryo-EM maps used in the benchmark dataset. The second column, “PDB-ID”, provides the PDB IDs of the structures associated with each map. The third column, “Resolution (Å)”, indicates the reported resolution of the deposited cryo-EM map. The fourth column, “Nchain”, specifies the number of chains in the corresponding PDB structure. The fifth column, “Nresidues”, shows the total number of residues in the PDB structure.

**Supplementary Table S5. AlphaFold2 single-chain structures for the high-resolution map dataset.** The “EMD-ID” and “PDB-ID” columns represent the EMDB and PDB identifiers for each entry, respectively. The “Chain_ID” column specifies the chain identifiers within the corresponding PDB structure. The “Uniprot_ID” column provides the UniProt ID linked to the single-chain protein entry in the AlphaFold database. For cases marked as “AFDB search”, we reported the entry ID from the AlphaFold Database. The “TM-Score” column reports the TM-Score metric, which measures the structural similarity between the AlphaFold2 model and the native structure.

**Supplementary Table S6. Modeling performance on the High resolution map dataset.** Each row represents a cryo-EM map identified by its EMD-ID. We reported seven metrics (Sequence Coverage, CC_box, F1 Score, CC_mask, Sequence Match, Backbone Precision, and Backbone Recall). We analyzed five methods, DMcloud, DiffModeler, EMBuild, DEMO-EM2, and ModelAngelo across multiple evaluation metrics. Each method’s performance is reported under the respective column for each metric. For reference, “AF2 Model Superimposition” represents models obtained by directly superimposing AlphaFold Database models onto the reference PDB structures. The blank data corresponds to the method that failed to build the model for the target.

**Supplementary Table S7. The DNA/RNA-protein complex benchmark dataset.** The first column, “EMD-ID”, lists the EMDB IDs of the cryo-EM maps used in the benchmark dataset. The second column, “PDB-ID”, provides the PDB IDs of the structures associated with each map. The third column, “Resolution (Å)”, indicates the reported resolution of the deposited cryo-EM map. The fourth and fifth columns (“drna_chains” and “drna_residues”) specify the number of nucleotide chains and the total number of nucleotide residues, respectively. The sixth and seventh columns (“protein_chains” and “protein_residues”) show the number of protein chains and the total number of amino acid residues in each PDB structure.

**Supplementary Table S8. Template structure analysis for the DNA/RNA-protein complex dataset.** The “EMD-ID” and “PDB-ID” columns represent the EMDB and PDB identifiers for each entry, respectively. The “Chain_ID” column specifies the chain identifiers within the corresponding PDB structure. The “Template“ column provides the PDB ID with the chain ID used as the template structure. The “TM-Score” column reports the TM-Score metric, which measures the structural similarity between the template and the native structure. To compute the TM-score for DNA/RNA structure, we used USalign. For the protein region, the “Uniprot_ID” column provides the UniProt ID linked to the single-chain protein entry in the AlphaFold database.

**Supplementary Table S9. Modeling performance of Methods on DNA/RNA-protein complex dataset** Each row represents a specific cryo-EM map identified by its EMD-ID. We reported five metrics (Sequence Coverage, Backbone F1 Score, Sequence Match, Backbone Precision, and Backbone Recall) for DNA/RNA and protein regions, respectively. We also report CC_box and CC_mask for whole DNA/RNA-protein complex models. For DNA/RNA regions, We analyzed four methods, DMcloud, VESPER, CryoREAD, and ModelAngelo, across multiple evaluation metrics. For protein regions, we analyzed DMcloud, DiffModeler and ModelAngelo. Each method’s performance is reported under the respective column for each metric. The blank data corresponds to the method that failed to build the model for the target.

**Supplementary Table S10. Modeling Performance of DMcloud on six large complex targets** The first column, “EMD-ID”, lists the EMDB IDs of the cryo-EM maps. The second column, “PDB-ID”, provides the PDB IDs of the structures associated with each map. The third column, “Resolution (Å)”, indicates the reported resolution of the deposited cryo-EM map. The fourth column, “Nchain”, specifies the number of chains in the corresponding PDB structure. The fifth column, “Nresidues”, shows the total number of residues in the PDB structure. We reported seven metrics (Sequence Coverage, CC_box, F1 Score, CC_mask, Sequence Match, Backbone Precision, and Backbone Recall).

**Supplementary Table S11. The computational time of DMcloud** The “EMD-ID” and “PDB-ID” columns represent the EMDB and PDB identifiers for each entry, respectively. The third column, ”Resolution (Å)”, indicates the reported resolution of the deposited cryo-EM map. The fourth column, “Nchain”, specifies the number of nucleotide chains in the corresponding PDB structure. The fifth column, “Nresidues”, shows the total number of amino acids or nucleotide residues in the PDB structure. The sixth column, “backbone detection (hours)” and “phosphate & sugar detection (hours)” shows the computational time (hours) of the diffusion model and 3D-CNN, respectively. The seventh column “DMcloud” shows the computational time (hours) of full pipeline of the DMcloud protocol.

## Acknowledgments

This work was partly supported by the National Institutes of Health (R01GM133840, R35GM158267, R21AI187928) to DK and the National Science Foundation (DBI2433490 to GT, IIS2211598, DBI2146026, and DBI2422620 to DK).

## Code Availability

DMcloud is available as a web server at https://em.kiharalab.org/algorithm/DMcloud. Computed models by DMcloud are available at https://zenodo.org/records/17458717.

## Author Contributions

GT and DK conceived the study. GT designed and implemented DMcloud and computed the results. XW and HZ participated in implementing the diffusion model part. HZ computed the part of the results. YZ prepared DNA/RNA datasets. JHP developed the DMcloud web server. DK and GT analyzed the results. GT drafted the manuscript and DK edited it. All the authors read and approved the manuscript.

---

**Extended Data Figure 1.**
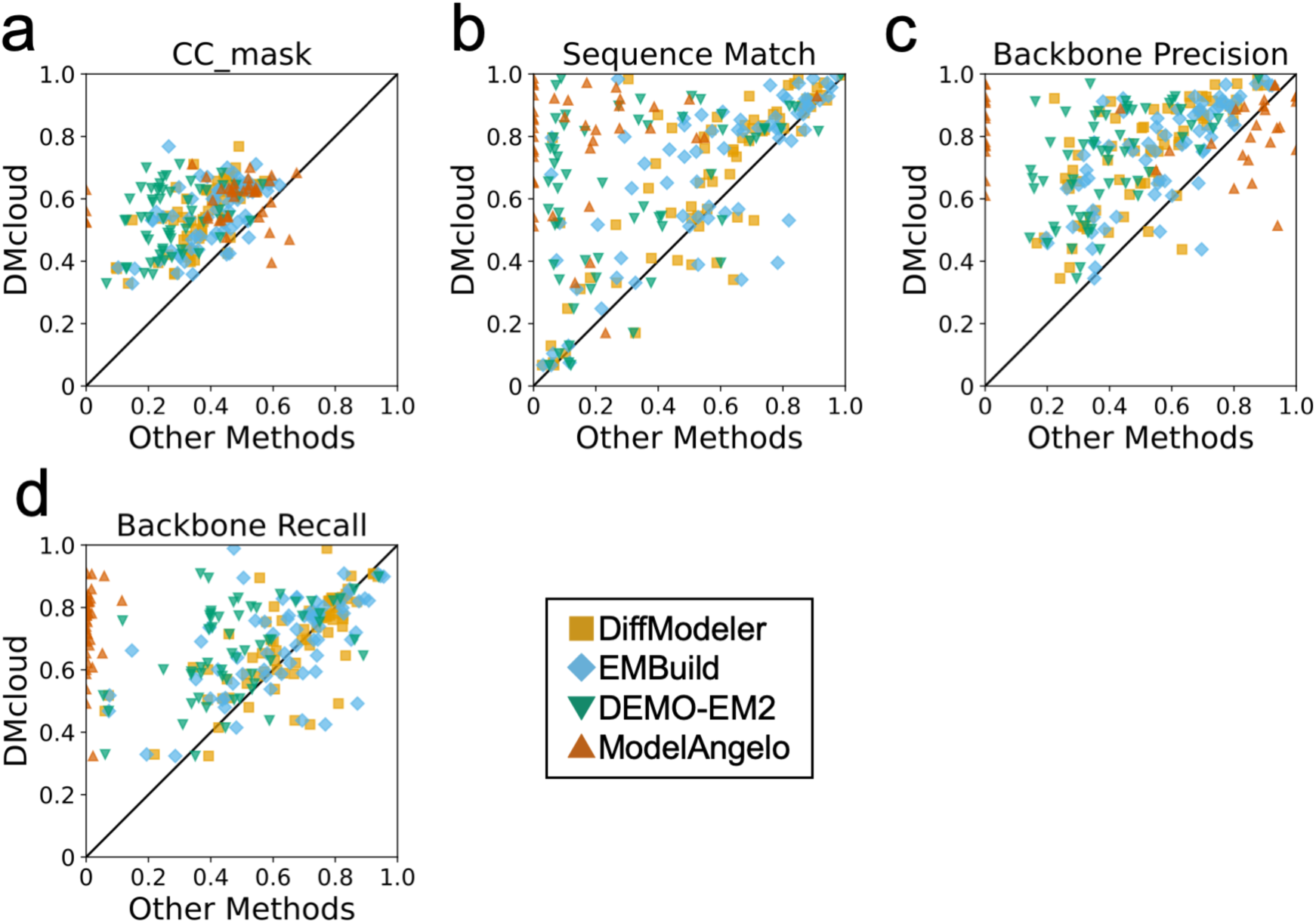
Evaluations of models for the intermediate-resolution dataset using four additional metrics. Each plot compares the performance of DMcloud (y-axis) with four other methods (x-axis), DiffModeler, EMBuild, DEMO-EM2, and ModelAngelo. **a.** CC_mask, local fit within the modeled region; **b.** Sequence Match, the fraction of correct residue type and alignment; **c.** Backbone Precision, accuracy of backbone atom position in the predicted model; **d.** Backbone Recall, completeness of backbone coverage in the reference PDB structure. Source data is provided in Supplementary Table S3.

**Extended Data Figure 2.**
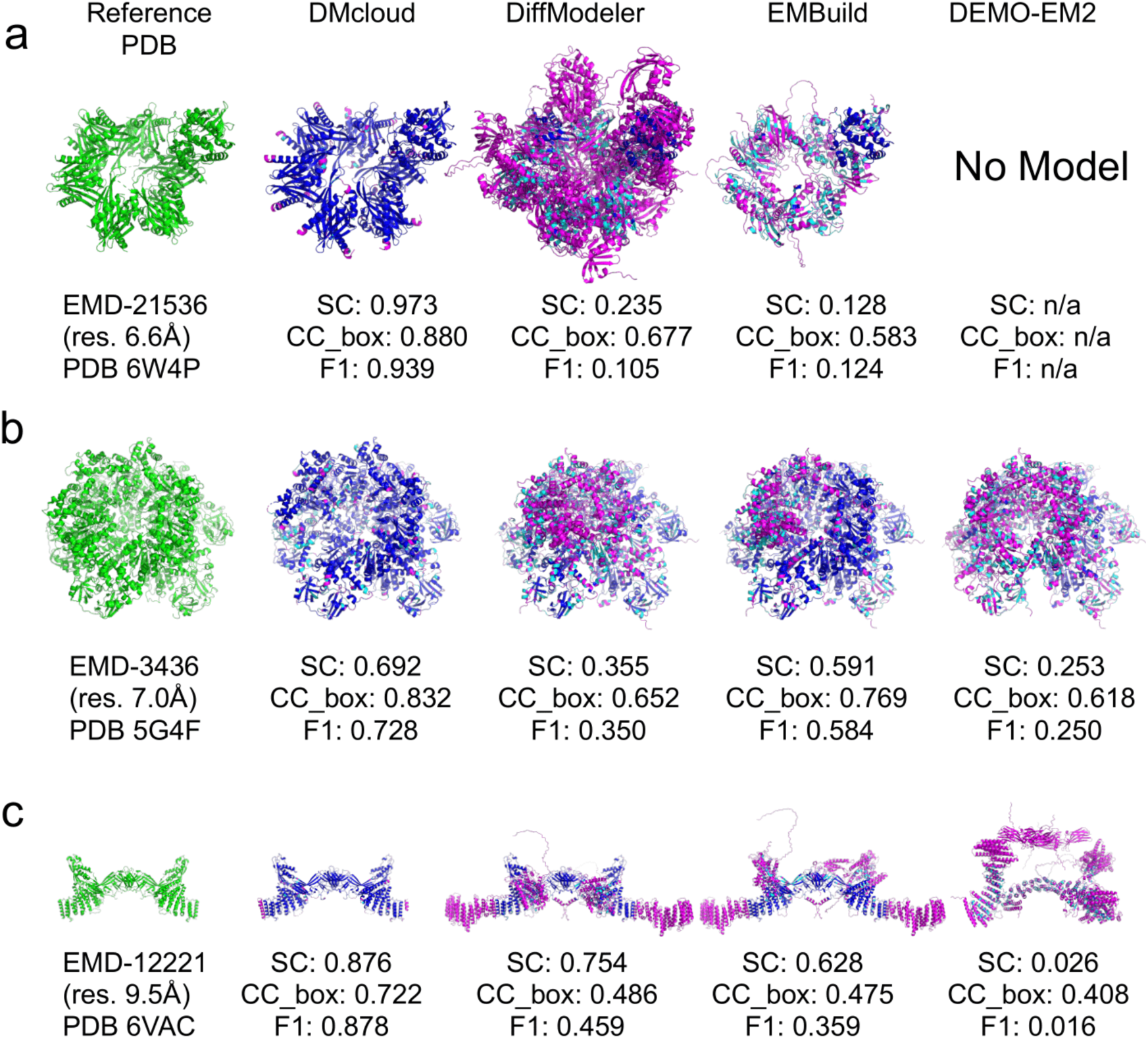
Examples of modeling results of DMcloud and the other three methods on three cryo-EM maps at intermediate resolutions. Each row corresponds to a different target: **a.** EMD-21536 (res. 6.6 Å, PDB 6W4P), **b.** EMD-3436 (res. 7.0 Å, PDB 5G4F), and **c.** EMD-12221 (res. 9.5 Å, PDB 6VAC). The first column shows the reference PDB structure, followed by models predicted by DMcloud, DiffModeler, EMBuild, and DEMO-EM2. Results of ModelAngelo was not shown in this figure because a ModelAngelo could not generate models at all for EMD-21536, and for EMD-3436 and EMD-12221, ModelAngelo generated fragmented structures with SC: 0.00. For EMD-21536, DEMO-EM2 could not generate any models. The models are color-coded in the same way as in Figure 2: magenta representing regions where the predicted position deviates more than 3 Å from the reference PDB structure, cyan indicating correct positional prediction but incorrect amino acid types, and blue showing correctly predicted amino acids within 3 Å of the reference PDB structure. Sequence Coverage (SC), CC_box, and F1 score (F1) values are shown for each model.

**Extended Data Figure 3.**
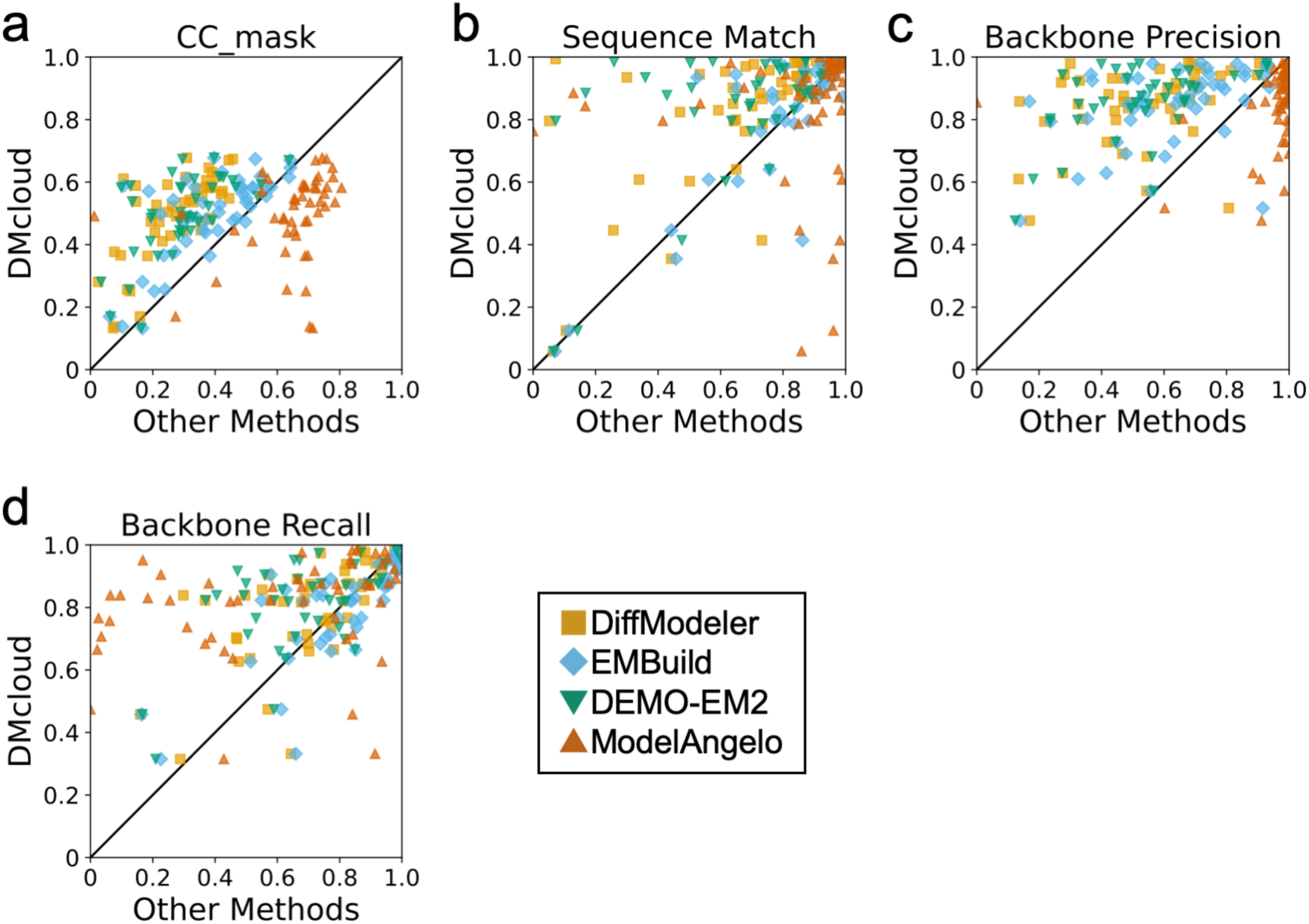
Evaluations of models for the high-resolution dataset using four additional metrics. Each plot compares the performance of DMcloud (y-axis) with four other methods (x-axis), DiffModeler, EMBuild, DEMO-EM2, and ModelAngelo. **a.** CC_mask, **b.** Sequence Match, **c.** Backbone Precision, **d.** Backbone Recall. Source data is provided in Supplementary Table S6.

**Extended Data Figure 4.**
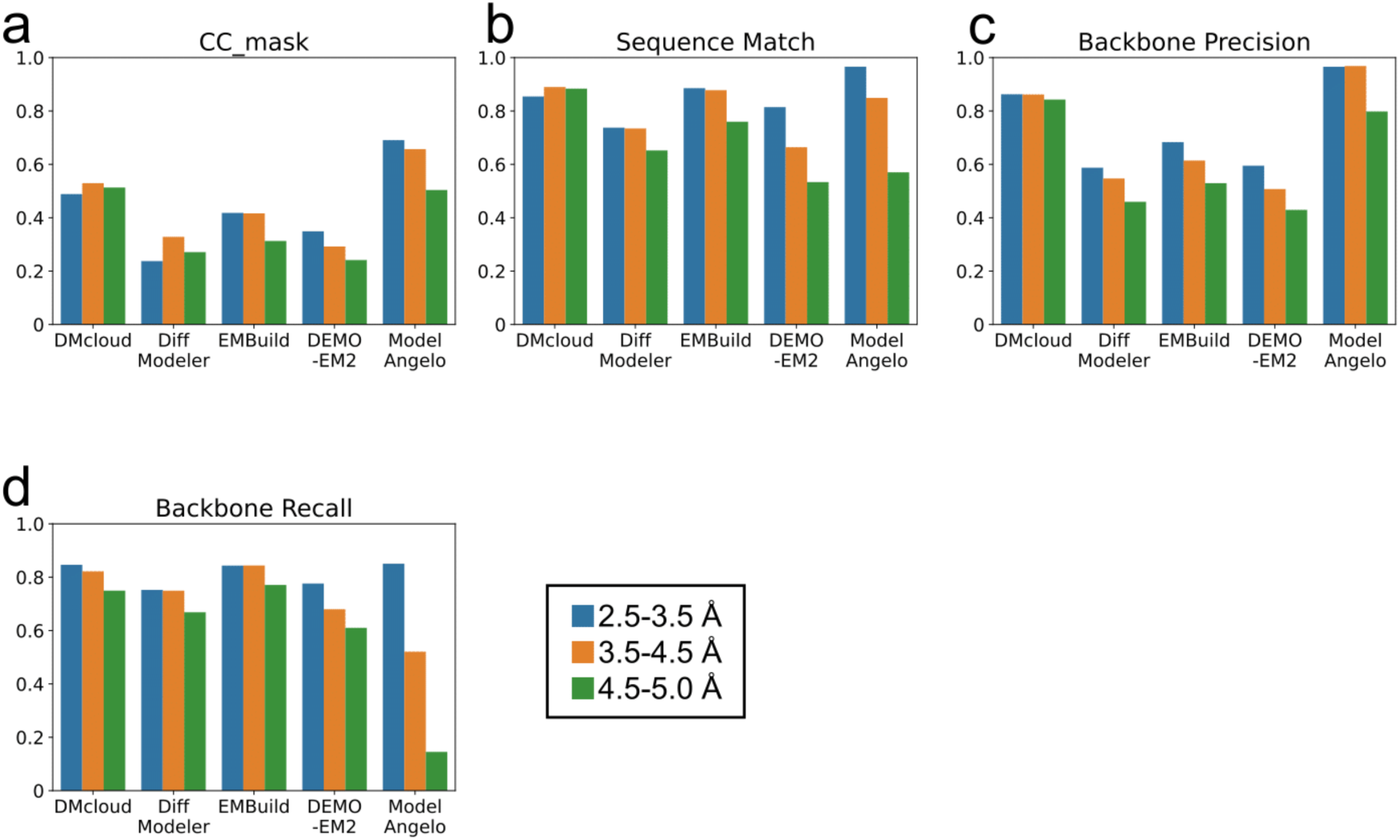
Evaluations of models for the high-resolution dataset using four additional metrics at different resolution ranges. Each plot shows the performance of DMcloud, DiffModeler, EMBuild, DEMO-EM2 and ModelAngelo across three different map resolution ranges. Each colored bar represents a different resolution range: 2.5-3.5 Å (blue), 3.5-4.5 Å (orange), and 4.5-5.0 Å (green). **a.** CC_mask, **b.** Sequence Match, **c.** Backbone Precision, **d.** Backbone Recall.

**Extended Data Figure 5.**
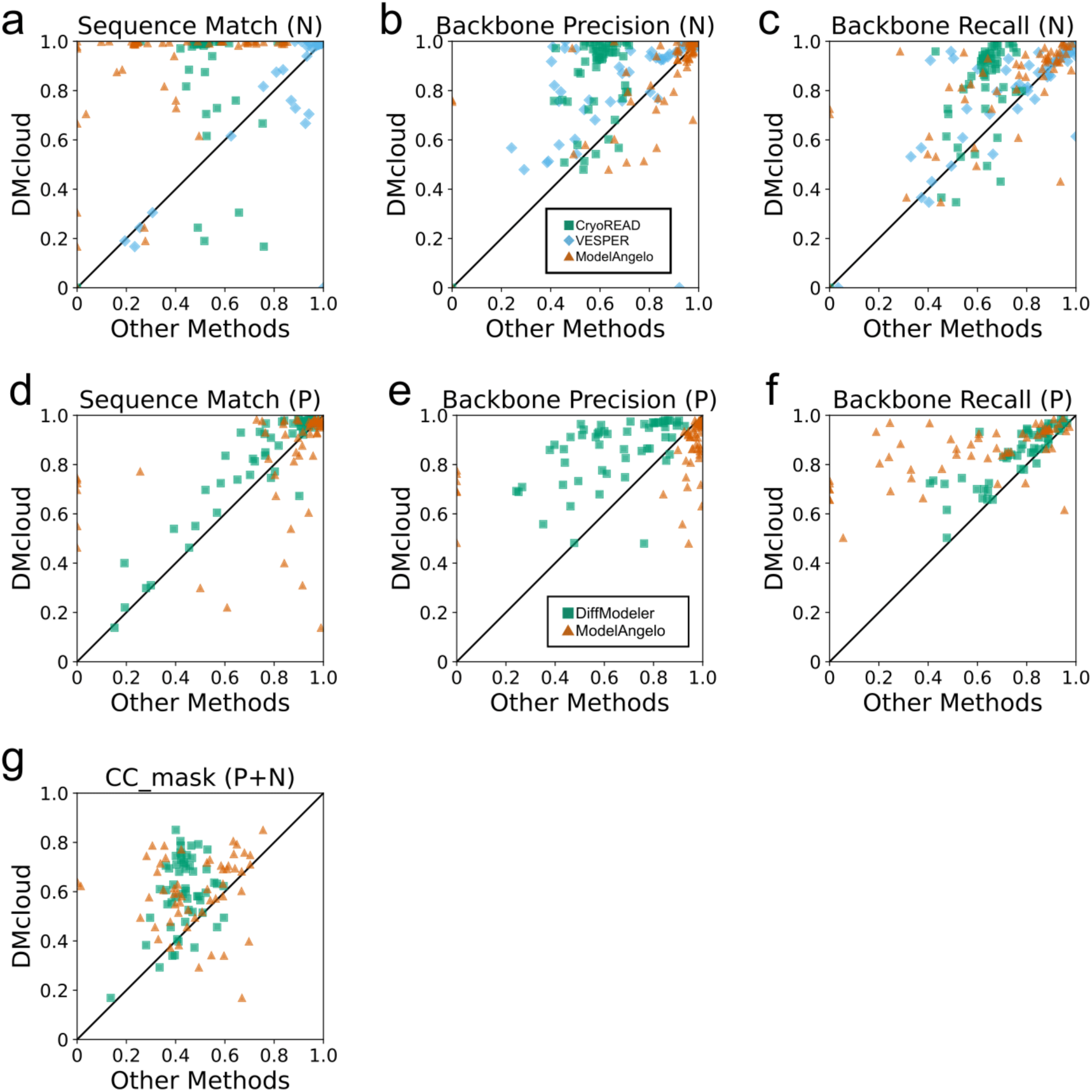
Evaluations of models for the DNA/RNA-protein complex dataset using seven additional metrics. Each plot compares the performance of DMcloud (y-axis) with three other methods (x-axis) **a–c**, Comparison with CryoREAD, VESPER, and ModelAngelo for DNA/RNA regions: **a.** CC_mask, **b.** Sequence Match, and **c.** Backbone Precision. **d–f**, Comparison with DiffModeler and ModelAngelo for protein regions: **d.** Backbone Recall, **e.** Backbone precision, and **f.** Backbone F1-score. **g**. Comparison for the entire DNA/RNA–protein complex using CC_mask.

**Extended Data Figure 6.**
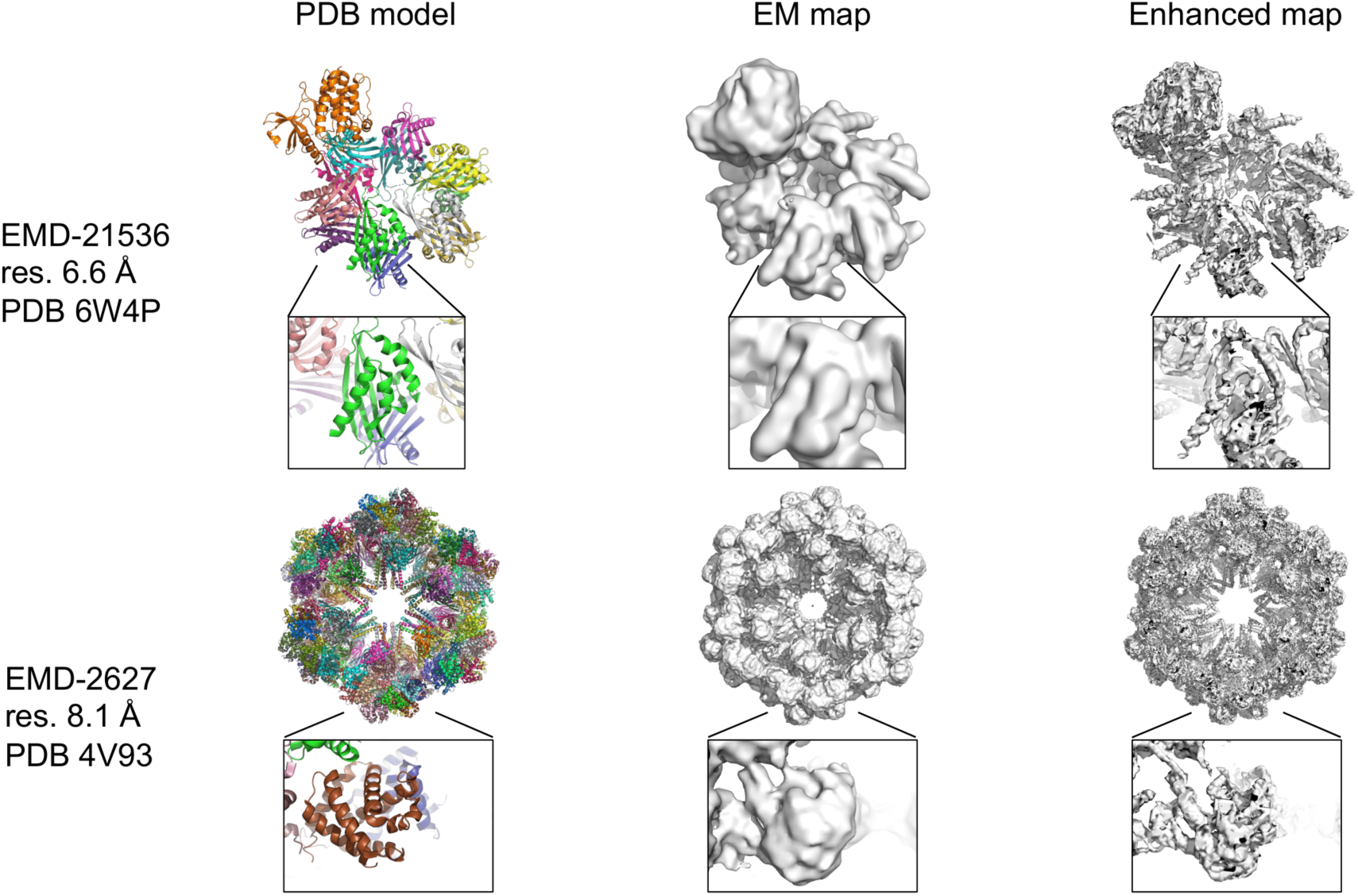
Examples of enhanced maps computed by the diffusion model. Two examples are shown that highlight the effect of the diffusion model on cryo-EM maps. EMD-21536 (resolution: 6.6 Å, PDB: 6W4P) and EMD-2627 (resolution: 8.1 Å, PDB: 4V93). The left column shows the atomic model from the corresponding PDB entry. The middle column shows the original cryo-EM map visualized at the author-recommended contour level. The right column displays the enhanced map generated by our diffusion model. We used a surface threshold of 1.0. Insets highlight local improvements in the enhanced maps.

**Extended Data Figure 7.**
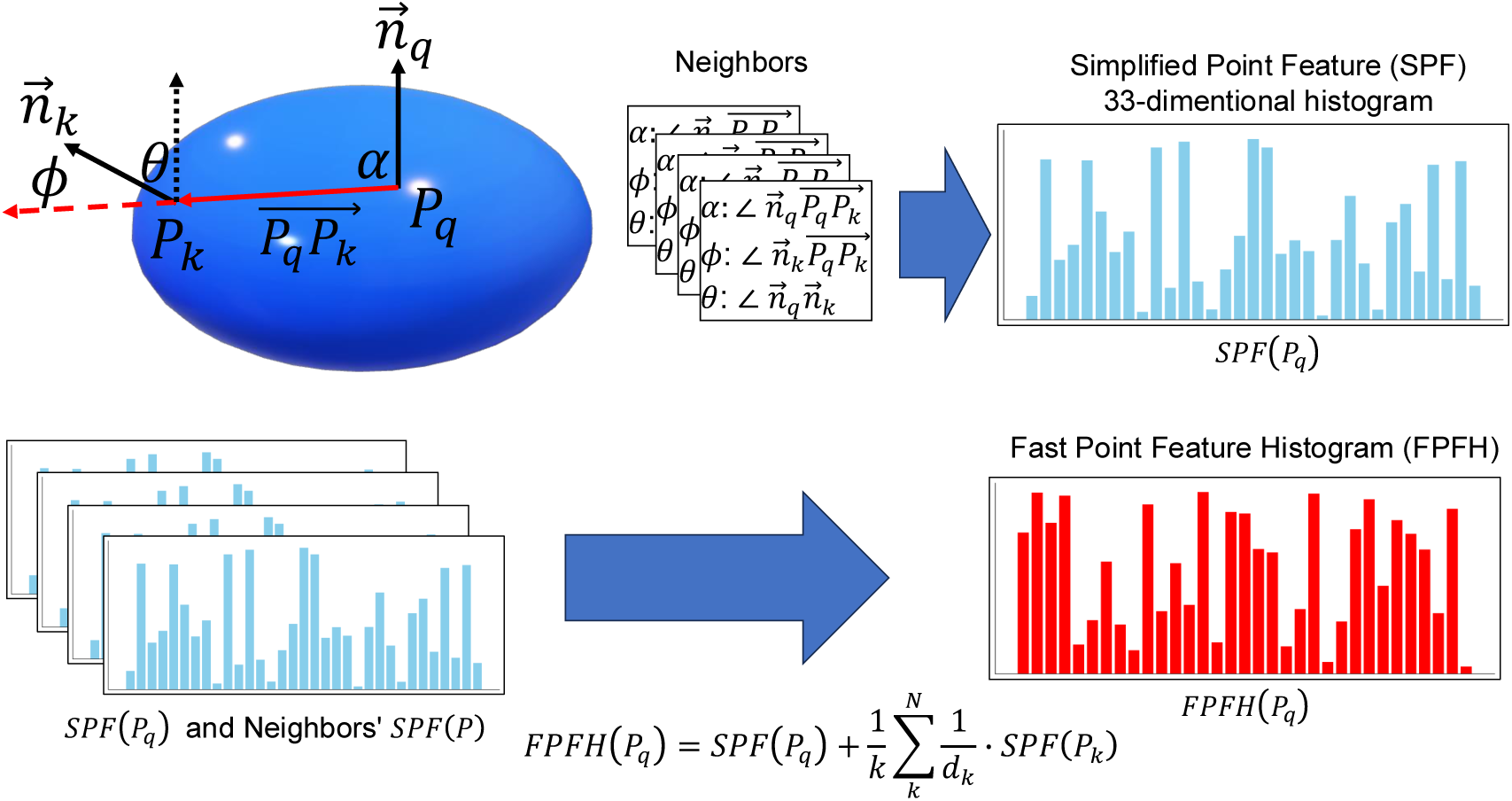
Overview of the Fast Point Feature Histogram (FPFH) This figure illustrates the process of computing the Fast Point Feature Histogram (FPFH) for a query point *P_q_* in a 3D point cloud. The FPFH is computed by first identifying neighboring points *P_k_* within a specified radius. For each point, normal vector is computed by fitting a plane to the local neighbor points. In addition, three angular features (*α*, *φ*, and *θ*) are computed between the query point *P_q_* and its neighbor point *P_k_*. *α* is the angle between the normal vector (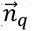) at the *P_q_* and the vector (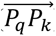) from *P_q_* to *P_k_*. *φ* is the angle between the normal vector (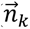) at the neighbor point *P_k_* and the vector 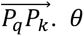 is the angle between 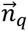 and 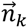. These angle values are summarized into 33-dimentional histogram (Simplified Point Feature, SPF) for the query point. Finally, FPFH for the query point is computed by combining the SPF of the query point and the weighted average of the SPFs of its neighbors. The weight is defined by the inverse of the distance (*d_k_*) between *P_q_* and *P_k_*.

**Extended Data Figure 8.**
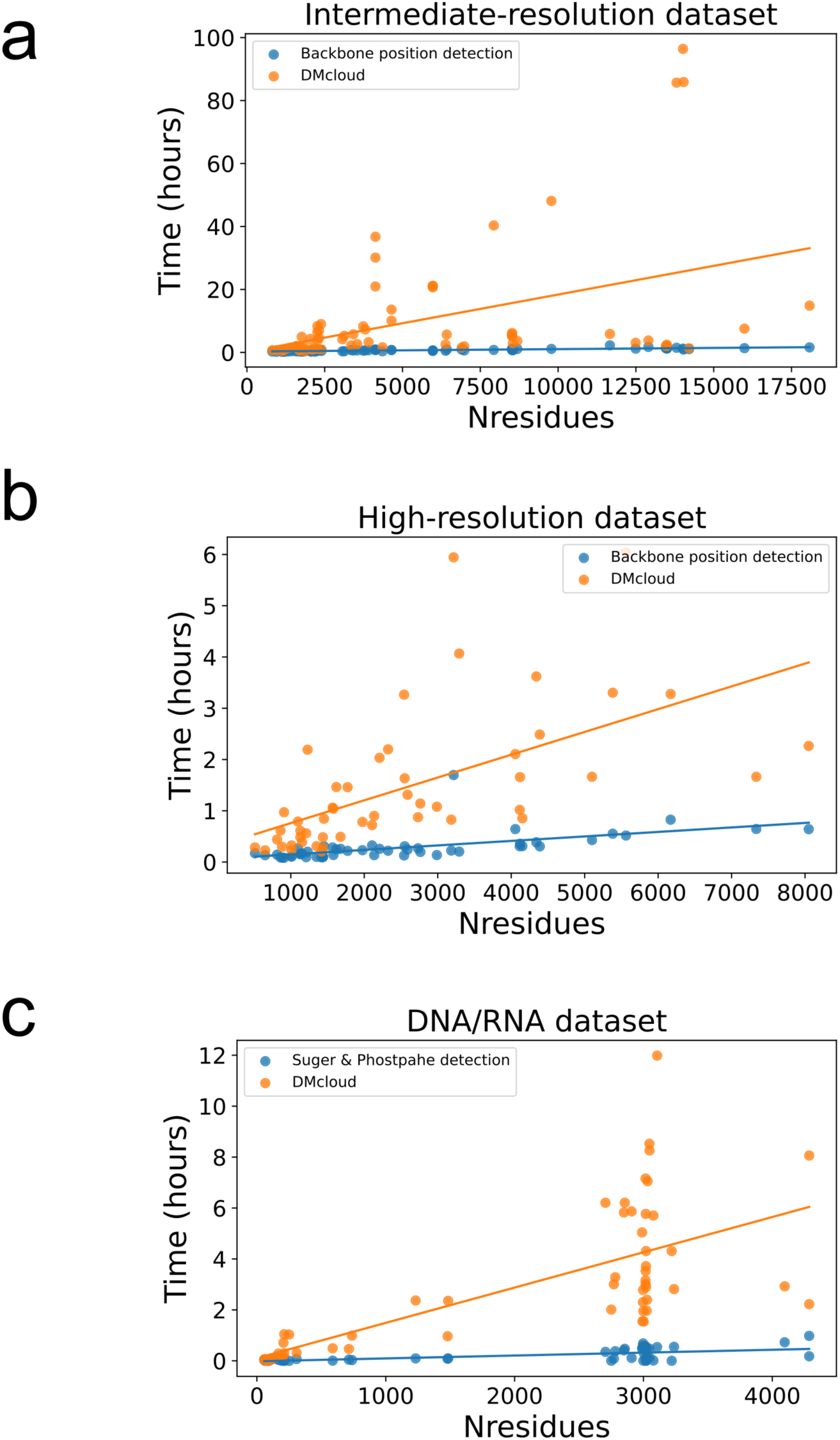
The computational time of DMcloud. Computational time of the DMcloud protocol is shown for **a.** the intermediate-resolution dataset (71 targets), **b.** the high-resolution dataset (50 targets), and **c.** the DNA/RNA-protein complex dataset (55 targets). We used a single GPU card (Nvidia RTX A6000 with 48 GB memory) and a CPU (Intel Xeon E5-1650 with 12 cores). In panels a and b, blue plots represent the computational time for backbone position detection by the diffusion model, while orange plots indicate the total computational time of DMcloud, including backbone position detection, point-cloud matching, and model assembly. In panel c, blue plots show the computational time for phosphate and sugar position detection using the 3D-CNN from CryoREAD, while orange plots represent the total computational time of DMcloud for DNA/RNA modeling. The source data are provided in Supplementary Table S11.

## References

1 Yip, K. M., Fischer, N., Paknia, E., Chari, A. & Stark, H. Atomic-resolution protein structure determination by cryo-EM. Nature 587, 157–161, doi:10.1038/s41586-020-2833-4 (2020).

2 Murata, K. & Wolf, M. Cryo-electron microscopy for structural analysis of dynamic biological macromolecules. Biochim Biophys Acta Gen Subj 1862, 324–334, doi:10.1016/j.bbagen.2017.07.020 (2018).

3 Farheen, F., Terashi, G., Zhu, H. & Kihara, D. AI-based methods for biomolecular structure modeling for Cryo-EM. Curr Opin Struct Biol 90, 102989, doi:10.1016/j.sbi.2025.102989 (2025).

4 Nakamura, T., Wang, X., Terashi, G. & Kihara, D. DAQ-Score Database: assessment of map-model compatibility for protein structure models from cryo-EM maps. Nat Methods 20, 775–776, doi:10.1038/s41592-023-01876-1 (2023).

5 Zhu, H., Terashi, G., Farheen, F., Nakamura, T. & Kihara, D. AI-based quality assessment methods for protein structure models from cryo-EM. Curr Res Struct Biol 9, 100164, doi:10.1016/j.crstbi.2025.100164 (2025).

6 Terashi, G., Wang, X., Prasad, D., Nakamura, T. & Kihara, D. DeepMainmast: integrated protocol of protein structure modeling for cryo-EM with deep learning and structure prediction. Nat Methods 21, 122–131, doi:10.1038/s41592-023-02099-0 (2024).

7 Li, S., Terashi, G., Zhang, Z. & Kihara, D. Advancing structure modeling from cryo-EM maps with deep learning. Biochem Soc Trans 53, 259–265, doi:10.1042/BST20240784 (2025).

8 Berman, H. M. et al. The Protein Data Bank. Nucleic Acids Res 28, 235–242, doi:10.1093/nar/28.1.235 (2000).

9 Jumper, J. et al. Highly accurate protein structure prediction with AlphaFold. Nature 596, 583–589, doi:10.1038/s41586-021-03819-2 (2021).

10 Han, X., Terashi, G., Christoffer, C., Chen, S. & Kihara, D. VESPER: global and local cryo-EM map alignment using local density vectors. Nat Commun 12, 2090, doi:10.1038/s41467-021-22401-y (2021).

11 Pettersen, E. F. et al. UCSF Chimera--a visualization system for exploratory research and analysis. J Comput Chem 25, 1605–1612, doi:10.1002/jcc.20084 (2004).

12 Chacon, P. & Wriggers, W. Multi-resolution contour-based fitting of macromolecular structures. J Mol Biol 317, 375–384, doi:10.1006/jmbi.2002.5438 (2002).

13 Kawabata, T. Multiple subunit fitting into a low-resolution density map of a macromolecular complex using a gaussian mixture model. Biophys J 95, 4643–4658, doi:10.1529/biophysj.108.137125 (2008).

14 Zhang, Z. et al. DEMO-EM2: assembling protein complex structures from cryo-EM maps through intertwined chain and domain fitting. Brief Bioinform 25, doi:10.1093/bib/bbae113 (2024).

15 Frigo, M. & Johnson, S. G. The design and implementation of FFTW3. Proceedings of the IEEE 93, 216–231 (2005).

16 He, J., Lin, P., Chen, J., Cao, H. & Huang, S. Y. Model building of protein complexes from intermediate-resolution cryo-EM maps with deep learning-guided automatic assembly. Nat Commun 13, 4066, doi:10.1038/s41467-022-31748-9 (2022).

17 Wang, X., Zhu, H., Terashi, G., Taluja, M. & Kihara, D. DiffModeler: large macromolecular structure modeling for cryo-EM maps using a diffusion model. Nat Methods 21, 2307–2317, doi:10.1038/s41592-024-02479-0 (2024).

18 Ho, J., Jain, A. & Abbeel, P. Denoising diffusion probabilistic models. Advances in neural information processing systems 33, 6840–6851 (2020).

19 Zhu, W., Shenoy, A., Kundrotas, P. & Elofsson, A. Evaluation of AlphaFold-Multimer prediction on multi-chain protein complexes. Bioinformatics 39, doi:10.1093/bioinformatics/btad424 (2023).

20 Fischler, M. A. & Bolles, R. C. Random sample consensus: a paradigm for model fitting with applications to image analysis and automated cartography. Communications of the ACM 24, 381–395 (1981).

21 Ester, M., Kriegel, H.-P., Sander, J. & Xu, X. in kdd. 226-231.

22 Jamali, K. et al. Automated model building and protein identification in cryo-EM maps. Nature 628, 450–457, doi:10.1038/s41586-024-07215-4 (2024).

23 Terashi, G. & Kihara, D. De novo main-chain modeling for EM maps using MAINMAST. Nat Commun 9, 1618, doi:10.1038/s41467-018-04053-7 (2018).

24 Rusinkiewicz, S. & Levoy, M. in Proceedings third international conference on 3-D digital imaging and modeling. 145–152 (IEEE).

25 Perron, L. in International Conference on Principles and Practice of Constraint Programming. 2-2 (Springer).

26 Huang, R. et al. Unfolding the mechanism of the AAA+ unfoldase VAT by a combined cryo-EM, solution NMR study. Proc Natl Acad Sci U S A 113, E4190–4199, doi:10.1073/pnas.1603980113 (2016).

27. Song, J., Meng, C. & Ermon, S. Denoising diffusion implicit models. arXiv preprint arXiv:2010.02502 (2020).

28 Varadi, M. et al. AlphaFold Protein Structure Database: massively expanding the structural coverage of protein-sequence space with high-accuracy models. Nucleic Acids Res 50, D439–D444, doi:10.1093/nar/gkab1061 (2022).

29 Rusu, R. B., Blodow, N. & Beetz, M. in 2009 IEEE international conference on robotics and automation. 3212–3217 (IEEE).

30 Perron, L. in Principles and Practice of Constraint Programming–CP 2011: 17th International Conference, CP 2011, Perugia, Italy, September 12-16, 2011. Proceedings 17. 2-2 (Springer).

31 Mariani, V., Biasini, M., Barbato, A. & Schwede, T. lDDT: a local superposition-free score for comparing protein structures and models using distance difference tests. Bioinformatics 29, 2722–2728, doi:10.1093/bioinformatics/btt473 (2013).

32 Wang, X., Terashi, G. & Kihara, D. CryoREAD: de novo structure modeling for nucleic acids in cryo-EM maps using deep learning. Nat Methods 20, 1739–1747, doi:10.1038/s41592-023-02032-5 (2023).

33 Liebschner, D. et al. Macromolecular structure determination using X-rays, neutrons and electrons: recent developments in Phenix. Acta Crystallogr D Struct Biol 75, 861–877, doi:10.1107/S2059798319011471 (2019).

34 Sloutsky, R. et al. Heterogeneity in human hippocampal CaMKII transcripts reveals allosteric hub-dependent regulation. Sci Signal 13, doi:10.1126/scisignal.aaz0240 (2020).

35 Leneva, N., Kovtun, O., Morado, D. R., Briggs, J. A. G. & Owen, D. J. Architecture and mechanism of metazoan retromer:SNX3 tubular coat assembly. Sci Adv 7, doi:10.1126/sciadv.abf8598 (2021).

36 Wang, R., Qi, X., Schmiege, P., Coutavas, E. & Li, X. Marked structural rearrangement of mannose 6-phosphate/IGF2 receptor at different pH environments. Sci Adv 6, eaaz1466, doi:10.1126/sciadv.aaz1466 (2020).

37 Bunduc, C. M. et al. Structure and dynamics of a mycobacterial type VII secretion system. Nature 593, 445–448, doi:10.1038/s41586-021-03517-z (2021).

38 Niu, Y., Suzuki, H., Hosford, C. J., Walz, T. & Chappie, J. S. Structural asymmetry governs the assembly and GTPase activity of McrBC restriction complexes. Nat Commun 11, 5907, doi:10.1038/s41467-020-19735-4 (2020).

39 Li, J., Wu, J., Hall, C., Bai, X. C. & Choi, E. Molecular basis for the role of disulfide-linked alphaCTs in the activation of insulin-like growth factor 1 receptor and insulin receptor. Elife 11, doi:10.7554/eLife.81286 (2022).

40 Altschul, S. F. et al. Gapped BLAST and PSI-BLAST: a new generation of protein database search programs. Nucleic Acids Res 25, 3389–3402, doi:10.1093/nar/25.17.3389 (1997).

41 Crowe-McAuliffe, C. et al. Structural Basis for Bacterial Ribosome-Associated Quality Control by RqcH and RqcP. Mol Cell 81, 115–126 e117, doi:10.1016/j.molcel.2020.11.002 (2021).

42 Chen, X. et al. Cutting antiparallel DNA strands in a single active site. Nat Struct Mol Biol 27, 119–126, doi:10.1038/s41594-019-0363-2 (2020).

43 Braxton, J. R., Shao, H., Tse, E., Gestwicki, J. E. & Southworth, D. R. Asymmetric apical domain states of mitochondrial Hsp60 coordinate substrate engagement and chaperonin assembly. Nat Struct Mol Biol 31, 1848–1858, doi:10.1038/s41594-024-01352-0 (2024).

44 Eisele, M. R. et al. Expanded Coverage of the 26S Proteasome Conformational Landscape Reveals Mechanisms of Peptidase Gating. Cell Rep 24, 1301–1315 e1305, doi:10.1016/j.celrep.2018.07.004 (2018).

45 Chen, W. T., Chen, Y. C., Liou, H. H. & Chao, C. Y. Structural basis for cooperative oxygen binding and bracelet-assisted assembly of Lumbricus terrestris hemoglobin. Sci Rep 5, 9494, doi:10.1038/srep09494 (2015).

46 Ranson, N. A. et al. ATP-bound states of GroEL captured by cryo-electron microscopy. Cell 107, 869–879, doi:10.1016/s0092-8674(01)00617-1 (2001).

47 von Kugelgen, A., Alva, V. & Bharat, T. A. M. Complete atomic structure of a native archaeal cell surface. Cell Rep 37, 110052, doi:10.1016/j.celrep.2021.110052 (2021).

48 Sun, S. Y. et al. Cryo-ET of Toxoplasma parasites gives subnanometer insight into tubulin-based structures. Proc Natl Acad Sci U S A 119, doi:10.1073/pnas.2111661119 (2022).

49 Zhou, Q.-Y., Park, J. & Koltun, V. Open3D: A modern library for 3D data processing. arXiv preprint arXiv:1801.09847 (2018).

50 Bekker, G. J. et al. Protein Data Bank Japan: Computational Resources for Analysis of Protein Structures. J Mol Biol 437, 169013, doi:10.1016/j.jmb.2025.169013 (2025).

51 Bekker, G. J. et al. Protein Data Bank Japan: Improved tools for sequence-oriented analysis of protein structures. Protein Sci 34, e70052, doi:10.1002/pro.70052 (2025).

